# The role of protein shape in multiphasic separation within condensates

**DOI:** 10.1101/2024.08.26.606306

**Authors:** Vikas Pandey, Tomohisa Hosokawa, Yasunori Hayashi, Hidetoshi Urakubo

**Affiliations:** Department of Biomedical Data Science, Fujita Health University School of Medicine, 1-98 Dengakugakubo, Kutsukake-cho, Toyoake 470-1192, Japan; International Center for Brain Science (ICBS), Fujita Health University, 1-98 Dengakugakubo, Kutsukake-cho, Toyoake 470-1192, Japan; National Institute of Physiological Sciences, National Institutes of Natural Sciences, 5-1 Myodaiji-Higashiyama, Okazaki 444-8787, Japan; Department of Pharmacology, Kyoto University Graduate School of Medicine, Kyoto 606-8501, Japan

**Keywords:** Liquid–liquid phase separation, CaMKII, Monte Carlo simulation, Synaptic plasticity

## Abstract

Liquid–liquid phase separation (LLPS) of biological macromolecules leads to the formation of various membraneless organelles. LLPS can not only form homogenous condensates but also multilayered and multiphase condensates, which can mediate complex cellular functions. However, the factors that determine the topological features of multiphase condensates are not fully understood. Herein, we focused on Ca^2+^/calmodulin-dependent protein kinase II (CaMKII), a major postsynaptic protein that undergoes various forms of LLPS with other postsynaptic proteins, and present a minimalistic computational model that reproduces these forms of LLPS, including a form of two-phase condensates, phase-in-phase (PIP) organization. Analyses of this model revealed that the competitive binding of two types of client proteins is required for the PIP formation. The PIP only formed when CaMKII had high valency and a short linker length. Such CaMKII proteins exhibited a low surface tension, a modular structure, and slow diffusion. These properties are consistent with the functions required by CaMKII to store information at the synaptic level. Thus, the computational modeling reveals new structure–function relationships for CaMKII as a synaptic memory unit.

## Introduction

Liquid–liquid phase separation (LLPS) is an emerging concept in biology where macromolecules spontaneously form condensates in aqueous solution, often occurring in nucleic acids and proteins with multivalent domain interactions or intrinsically disordered regions (IDR)^1, 2, 3^. LLPS leads to the formation of organelles that lack membranes, such as transcription and translational machineries, ribonucleic acid (RNA) granules, stress bodies, nucleoli, chromosomes, and postsynaptic density (PSD)^4, 5^. By concentrating and organizing molecules in a small domain, LLPS facilitates the reactions mediated by these organelles. The functions of the condensates can be further elaborated using more complex forms of LLPS. Two or more phases with different components can coexist in single condensate in various topological configurations such as phase-in-phase (PIP), partially engulfed, or phase-on-phase structures (Fig. 1a)^4, 6^. The topological structure of a multicomponent condensate is influenced by interfacial tensions^7, 8, 9^, competitive binding^9, 10^, and charge patterning in IDR^11^ and such characteristics help cells to organize cellular components in functionally related subdomains^12, 13^. For instance, a two-phase condensate with outer and inner phase layers can facilitate sequential reactions; a molecule in the diluted phase must pass through the outer phase before reaching the inner phase, thereby forcing the reaction mediated by the outer layer to occur before that in the inner core^14^. Furthermore, because of Ostwald ripening, wherein small condensates spontaneously merge into a larger condensate, small condensates in a microdomain rarely persist^15,16^. This phenomenon can be circumvented by wrapping high-surface tension small condensates with the outer phase exhibiting low surface tension^6, 16, 17^. However, the factors that determine the topology have not been fully theoretically accounted for. Given that attempts to target LLPS as a therapeutic approach have already begun^18, 19^, a thorough understanding of the mechanism governing multicomponent condensate is critical.

**Fig. 1.**
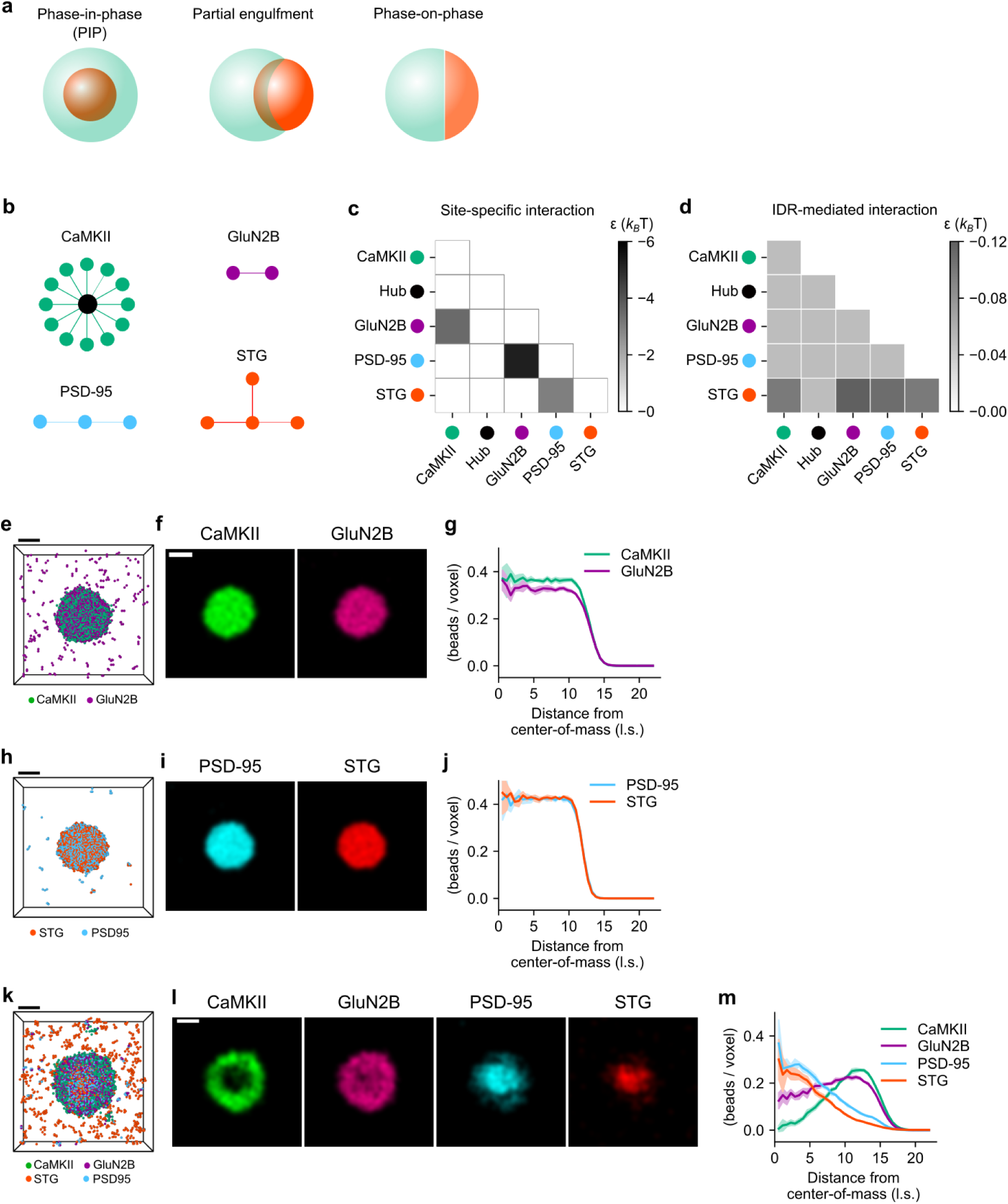
Various LLPS are reproduced in the Monte Carlo (MC) simulation of postsynaptic proteins. **a** Topological characteristics of the two-phase condensate. Depending on the surface and interfacial tensions, the two-phase condensate shows phase-in-phase (PIP) (left), partially engulfed (center), or even phase-on-phase structures^4, 6^. **b** To explore the multiphase condensates at postsynaptic spines, we modeled four synaptic proteins as a combination of beads and linkers. Filled circles denote beads, each of which occupied a lattice site, and lines denote linkers that connect two beads. **c** Interaction matrix for site-specific interactions. *ε* denotes binding energies (see Methods). **d** Interaction matrix for the intrinsically disordered region (IDR)-mediated interactions. **e–m** Protein distribution after the MC simulation. **e–g** Simulated condensate of the binary mixture of CaMKII and GluN2B. **e** Protein distribution in the three-dimensional reaction space. The front half was removed, and only the rear half is shown for visibility. **f** Normalized concentration levels at the section that divides the center of the condensate. **g** Radial distribution profiles (RDP) of protein concentrations from the center of the condensate. Means and standard deviations (SDs) were obtained based on data at five sampling times (Methods). **h–j** Simulated condensate of the binary mixture of CaMKII and GluN2B. **k–m** Simulated condensate of the mixture of all four proteins. Details for simulation and visualization are described in the Methods section. Scale bar, 10 l.s.

The factors which determine the efficacy with which a molecule undergoes LLPS have been explored using simulations, including Monte Carlo (MC) and Molecular Dynamics (MD) simulations^20, 21^. This approach enables efficient phase-plane analyses to evaluate the topological changes induced by continuous variables such as concentrations^22^. In line with this, researches targeting linear RNA-binding proteins have demonstrated that multiphase condensates are dependent on molecular concentrations due to the competitive binding between different molecular species^9, 10^. The effect of linkers length connecting each protein interaction domains has also been examined for a monomeric linear multivalent molecules^23, 24, 25, 26, 27^. However, molecules exhibiting LLPS are often not linear but multimeric nature, making them more complex overall^6, 28, 29^.

We targeted four postsynaptic proteins, CaMKII, *N*-methyl-D-aspartate (NMDA) receptor subunit GluN2B, PSD protein 95 (PSD-95), and α-amino-3-hydroxy-5-methyl-4-isoxazolepropionic acid receptor (AMPAR)-auxiliary subunit stargazin (STG). CaMKII has recently been shown to play a structural role at the synapse not just acting as a kinase mediating cellular signaling^30, 31^. This role is mediated by Ca^2+^/calmodulin-induced LLPS of the kinase domain of CaMKII with its binding partners, which include the carboxyl tail of GluN2B^5, 14^, through its rotationally symmetric dodeca- or tetradecameric structure^32^. The ternary mixture of STG–PSD-95–GluN2B forms a homogenous condensate^14, 33^. When CaMKII is present in the absence of Ca^2+^/calmodulin, CaMKII remains in the diluted phase, whereas STG–PSD-95–GluN2B still forms a homogenous condensate. However, when Ca^2+^/calmodulin is added to stimulate CaMKII, CaMKII organizes on the condensate as in PIP form; the CaMKII-GluN2B predominantly resides in the outer layer, whereas STG-PSD-95 occupies the inner core of this condensate^6, 9, 14, 34^. This structure persists even after the calcium is chelated by ethylene glycol tetraacetic acid (EGTA), consistent with the maintenance of long-term potentiation^32^. This spatial segregation facilitates the segregation of AMPA and NMDA receptors, thereby facilitating efficient neurotransmission^14^. Because the same set of proteins can assume different topological configurations of condensate depending on the conditions, we deemed that these proteins offer a versatile model for understanding the mechanism of the formation of multiphasic LLPS.

Therefore, we used the MC method on the Lattice-Based Structure and Stability Investigation (LaSSI)-simulation engine to model protein shapes and interactions and simulate their collective behaviors^6, 7, 8, 35, 36, 37, 38^. The model was built with a biologically plausible set of parameters that reproduced various experimentally observed LLPS^14, 33, 39, 40, 41^, and the simulated concentration dependence revealed that competitive binding to PSD-95 is required for the PIP formation. Through alterations in the CaMKII shape, we found that high valencies through the rotationally symmetric dodecameric structure and short linker length were both required for the PIP. The short linker length created a low surface tension, which is required to form the outer phase of the PIP structure while the high valency is required for the condensate formation itself. The simulation results also implied that the modular structure formation in the CaMKII–GluN2B condensate caused the slow diffusion of CaMKII. These results revealed new structure–function relationships for CaMKII as a synaptic memory unit. This is the first systematic and mechanistic study investigating the divergent structure of protein-regulated multiphase condensates.

## Results

### A minimal model of postsynaptic proteins reproduces the various forms of LLPS

To elucidate the mechanisms of condensate formation involving postsynaptic proteins, we used the LaSSI-simulation engine, which handles MC simulations in a three-dimensional (3D) lattice space where each lattice site is occupied by one bead, to simulate the collective behaviors of proteins^35^. Updates of bead locations are based on the Metropolis–Hastings algorithm that considers binding energies and spatial constraints, and its fast computation is advantageous for studying cellular-level LLPS^6, 7, 8, 35, 36, 37, 38^.

We first modeled the shapes of CaMKII, GluN2B, PSD-95, and STG (Fig. 1b). CaMKII was modeled as a combination of 12 binding beads representing the kinase domain and a hub bead representing the association domain (Fig. 1b, top left). Each pair of hub and binding beads was connected by a linker whose maximal length was 3 lattice site (l.s.), where l.s. denotes the unit length (see Methods). Those connections represented the hub-and-spoke architecture of CaMKII (Fig. 1b, top left). GluN2B was modeled as a pair of beads, mimicking the endogenous tetrameric receptor complex with GluN1 (Fig. 1b, top right)^14^. PSD-95 was modeled as a series of three beads, each representing a PDZ domain (Fig. 1b, bottom left). STG was modeled as T-shaped tetrameric beads representing the tetrameric, concentrated, and small-sized structure of STG^14^. The maximal linker lengths between the GluN2B, PSD-95, and STG beads were all set as 2 l.s., considering that their bead sizes should be similar to their linker lengths^14^. We did not map the lengths of these models to actual lengths as in many other schematic models^6, 7, 8, 35, 38^.

Herein, we modeled the site-specific interactions between the CaMKII binding beads and GluN2B, GluN2B and PSD-95, and PSD-95 and STG (Fig. 1c). The binding energies of these site-specific interactions^39, 42^ were determined based on previous experimental studies (see Methods). The site-specific interactions were modeled as the anisotropic interactions of LaSSI, as one-to-one (mutually exclusive) and anisotropic.

We further modeled IDR-mediated interactions. As shown in Fig. 1d, all constituent beads were set to be associated with two orders of magnitude weaker than the site-specific interactions (see Methods)^7, 8^. The carboxyl tails of GluN2B and STG, both of which were incorporated in *in vitro* experiments (Methods), are indeed IDRs, and the carboxyl tail of STG and PSD-95 is known to interact through their IDRs, which is attributed to the presence of arginine-rich segments^33^. These interactions were modeled as the isotropic interactions of LaSSI and were therefore one-to-many and isotropic. Other details, including the simulation time schedule, are described in the Methods section.

To validate the constructed model, we first simulated binary mixtures of CaMKII–GluN2B, PSD-95–STG, and GluN2B–PSD-95 (Fig. 1e–j, Supplementary Fig. 1a–c). Consistent with the preceding experimental and computational evidence^14, 33, 43^, they all showed homogeneous condensation (Fig. 1e–j, Supplementary Fig. 1a–c) and remained spherical as shown in their 3D profiles (Fig. 1e, h, Supplementary Fig. 1a). Their sectional images revealed the homogeneity of the condensate (Fig. 1f, i, Supplementary Fig. 1b), and the radial distribution profiles (RDP) from the center-of-mass showed a quantitative uniform distribution (Fig. 1g, j; Supplementary Fig. 1c). The ternary mixture of CaMKII–GluN2B–PSD-95 also underwent homogeneous condensation, which is again consistent with results from a previous study (Supplementary Fig. 1d–f)^44^.

The quaternary mixture of CaMKII, GluN2B, PSD-95, and STG underwent PIP condensation (Fig. 1k–m). CaMKII and GluN2B, located at the periphery of the condensate, formed the shell phase while PSD-95 and STG, concentrated at its center, formed the core phase (Fig. 1k–m). The core formation presumably came from the tight coupling of PSD-95 and STG, and GluN2B and its binding partner; CaMKII, was relegated to the periphery presumably because the CaMKII–GluN2B phase had a low surface tension^6, 7, 8, 9, 11, 34^. In contrast, when inactive CaMKII, GluN2B, PSD-95, and STG were mixed, the inactive CaMKII was dissipated in the diluted phase, and the latter three phases underwent homogeneous condensation (Supplementary Fig. 1g–i). Here, inactive CaMKII denotes the CaMKII devoid of the interactions, mimicking an inactive, closed conformation of the enzyme. Upon activation of CaMKII, the homogeneous condensate was transformed into a PIP condensate structure (Supplementary Video 1). This transition occurred autonomously, ensuring that PIP formation was independent of the initial spatial distribution of the proteins. Overall, our simulations successfully reproduced the various forms of phase-separated structures and their mesoscopic characteristics, thus validating the robustness of this model.

### Experimentally observed concentration dependence aligns with the computational model

We then tested the concentration dependency of the PIP organization in the model (Fig. 2 d–f, top). When we decreased the concentration of STG by three-fold or less, the condensate became homogenous. Similarly, when we increased the concentration of GluN2B and STG simultaneously by 1.5-fold or more, the condensate also became homogenous. Their topological shapes were clearly characterized in their rendered 3D shapes (Fig. 2 d–f, bottom left). To validate our simulation, we performed biochemical experiments using heterologously expressed and fluorescently labeled proteins (see Methods, Fig. 2a–c). The control set of concentrations (i.e., 20 µM CaMKII, 5 µM GluN2B, 15 µM PSD-95, and 20 µM STG) exhibited PIP organization (Fig. 2a). Conversely, a three-fold decrease in STG concentration to 5 µM produced homogenous condensates instead of PIP organization (Fig. 2b). Furthermore, a simultaneous two-fold increase of GluN2B and STG concentration (10 and 30 µM, respectively) produced a homogenous condensate (Fig. 2c). A small heterogeneity in the homogeneous condensate (Figs. 2e, f) was also consistent with the simulation (Fig. 2c). Overall, both simulation and biochemical experiments confirmed that PIP organization did not always appear but depended on the specific protein concentrations.

**Fig. 2.**
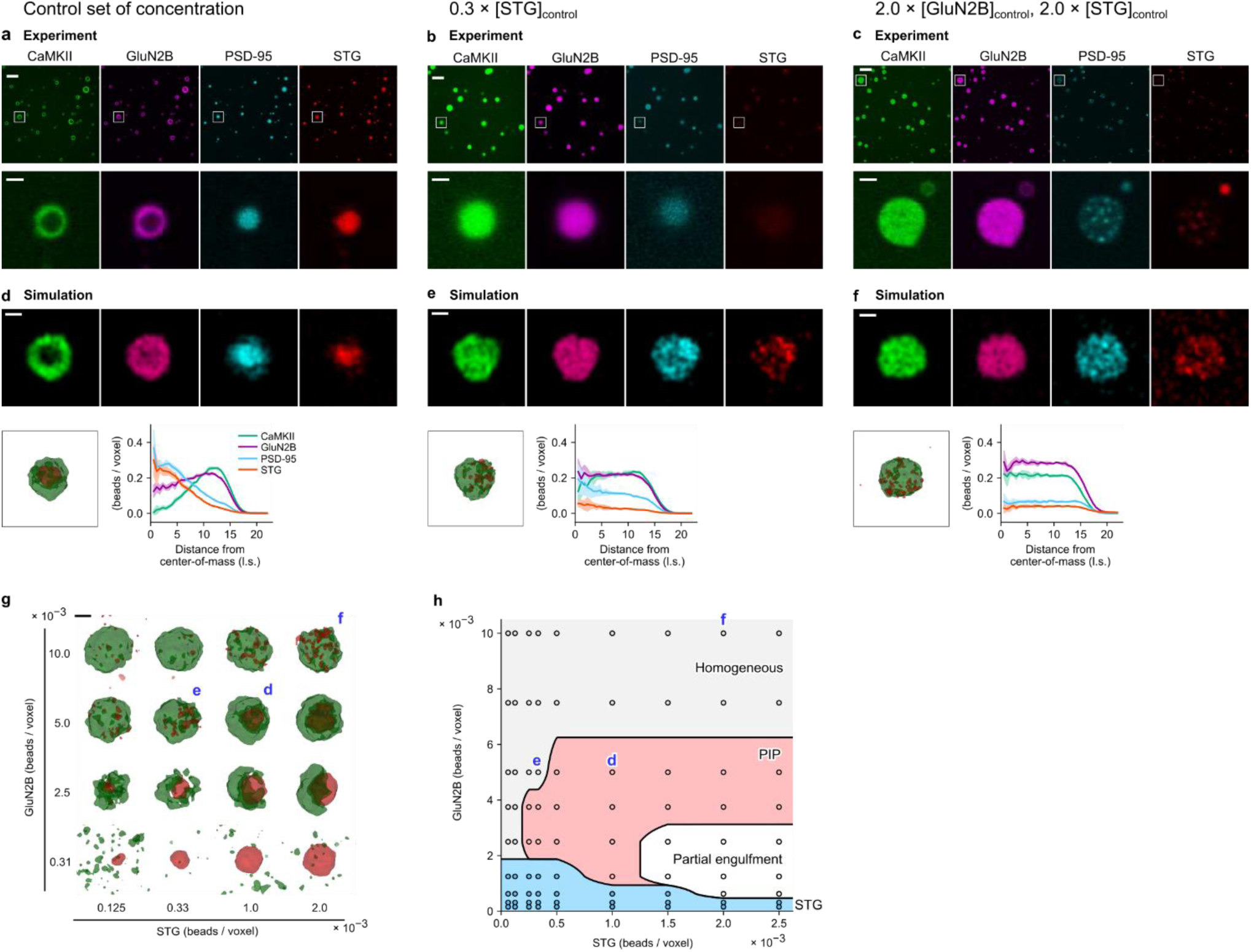
Either PIP or homogeneous condensate appears depending on the protein concentration. **a** Experimentally generated PIP structures with the quaternary mixture of postsynaptic proteins. Concentrations of CaMKII, GluN2B, PSD-95, and STG were 20, 5, 15, and 20 μM, respectively. Top row: fluorescent microscopic images of proteins. Each protein was labeled with a unique-colored fluorescent dye. Bottom row: high-magnification images of an example condensate. Scale bars, 5 and 1 μm for low- and high-magnification images, respectively. **b** Homogeneous condensate appears under the three-fold decrease in the STG concentration. **c** Homogeneous condensate appears under a simultaneous two-fold increase in concentrations of GluN2B and STG. **d–f** The replication of LLPS under the MC simulation. The control set of concentrations was 10, 5, 5, and 1 (in × 10^−3^ beads/voxel) for CaMKII, GluN2B, PSD-95, and STG, respectively. Top: Cross-sectional concentration levels of indicated proteins. Scale bar: 10 l.s. Bottom left: A rendered shape of CaMKII and STG phases. The CaMKII-enriched region is colored green while the STG-enriched region is colored red. Bottom right: RDPs. **g** Concentration-dependent shapes of LLPS. The cases (**d**), (**e**), and (**f**) are indicated. All condensates were rotated so that CaMKII-enriched regions are located on the left side. Scale bar: 10 l.s. **h** Phase diagram of the LLPS. Circles denote sampling points, and each colored region shares the same topological shape. Each region was determined based on visual inspection (see Supplementary Fig. 2). Sampling points (**d**), (**e**), and (**f**) are indicated.

Further simulation revealed the requirement of minimal GluN2B for the CaMKII–GluN2B condensation. At a GluN2B concentration below a threshold level, the CaMKII–GluN2B condensate did not appear and only the STG–PSD-95 condensate appeared (e.g., 0.31 × 10^−3^ beads/voxel in Fig. 2g, bottom; blue-shaded region in Fig. 2h). Similarly, if the concentration of GluN2B exceeded a certain level (gray-shaded region in Fig. 2h), this led to well-mixed homogeneous condensate comprising the four proteins (e.g., 10.0 × 10^−3^ beads/voxel in Fig. 2g). If the STG concentration was >1.5 × 10^−3^ beads/voxel, in some cases, the two-phase condensates formed a partial engulfment, but not a PIP (white region in Fig. 2h). Together, the two-phase condensates of the quaternary mixture only appeared in the appropriate concentration range of GluN2B and STG (red and white regions in Fig. 2h). As summarized in the phase diagram (Fig. 2h, Supplementary Fig. 2), each form clearly occupied its own concentration range, suggesting the existence of underlying mechanisms.

### PSD-95 sharing is required for PIP

We next examined the role of site-specific interaction between CaMKII and GluN2B. This interaction was important for the CaMKII–GluN2B phase of the PIP organization. The average number of GluN2B proteins bound to one CaMKII almost completely reflected the concentration of GluN2B (Fig. 3a) and corresponded to the volume of the CaMKII condensate (*R*^2^ = 0.97, Fig. 3b). Here, the CaMKII condensate was defined as the region of the largest volume among the group of regions where the CaMKII concentration exceeded the half maximal level (Methods).

**Fig. 3.**
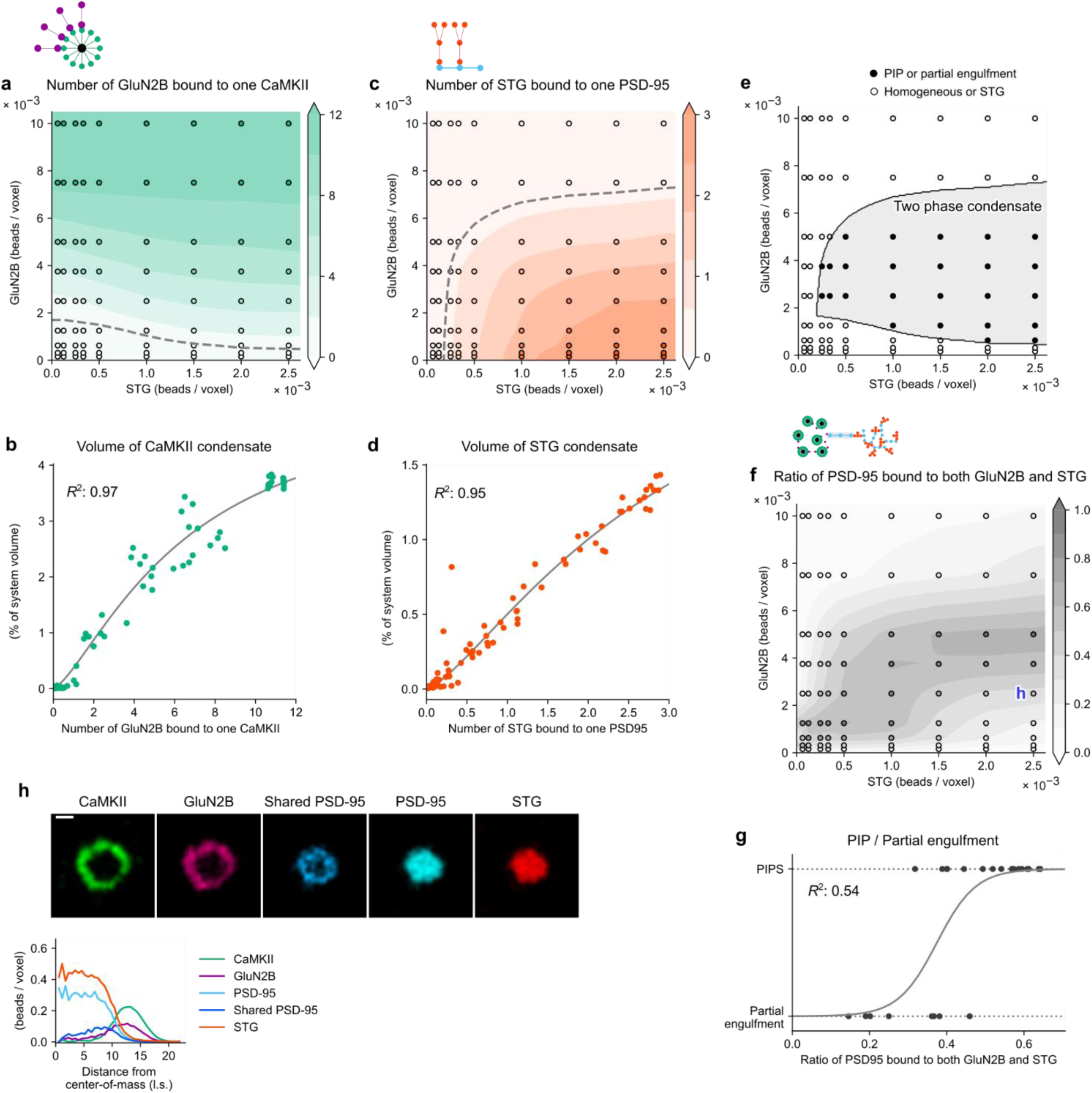
The concentration dependence of two-phase LLPS is explained by competitive binding. **a** Average numbers of GluN2B proteins bound to one CaMKII protein were plotted under the various concentration pairs of STG and GluN2B. Open circles indicate the observation while colored areas show their linear interpolation. **b** The GluN2B–CaMKII binding explains the volume of CaMKII condensate. Volumes of the largest CaMKII condensate were plotted against average numbers of GluN2B–CaMKII binding in the system. The Hill function was fitted, and the goodness-of-fit (*R*^2^) is shown. **c** The average number of STG proteins bound to one PSD-95 in the same concentration space of STG and GluN2B. **d** The STG–PSD-95 binding explains the volume of STG condensate. Volumes of the largest STG condensate were plotted against average numbers of STG–PSD-95 binding. The Hill function was fitted. **e** The two-phase separation is explained by the two types of binding. The gray-shaded area, which was derived from equivalued lines of average bindings in panels **a** and **c** (see gray-dashed lines), covered the two-phase separation with 100% accuracy. The procedure is shown in Supplementary Fig. 3. **f** Ratios of PSD-95 sharing by GluN2B and STG in condensates. **g** The PSD-95 sharing ratio in panel **f** discriminated the PIP and partial engulfment; shape data were taken from Fig. 2h. Efron’s *R*^2^ is displayed as a measure of the goodness-of-fit of the logistic function. **h** The sharing of PSD-95 appears at the interface region between phases of CaMKII and STG. The concentrations are indicated in panel **f**. For visibility, the RDP was generated only from the last time frame data. Scale bar: 10 l.s.

Similarly, we examined the site-specific interaction between STG and PSD-95, which was related to the STG–PSD-95 phase. As expected, the average number of STG proteins bound to one PSD-95 increased with increasing STG concentration (Fig. 3c), whereas the number of STG proteins bound to PSD-95 decreased with the increasing concentration of GluN2B (Fig. 3c). This is because GluN2B competes with STG with the same binding site on PSD-95. Consequently, the STG volume decreased in the high concentration range of GluN2B (Fig. 3d). The CaMKII and STG condensates were required for the two-phase condensate because the two-phase region (Fig. 2h) was determined by the binding of certain numbers of GluN2B–CaMKII molecules and STG–PSD-95 molecules (Fig. 3e and Supplementary Fig. 3).

Further, PIP and partial engulfment were discriminated by the average ratio of PSD-95 bound to both GluN2B and STG (Fig. 3f, g). The greater the number of PSD-95 bound to both GluN2B and STG, the more likely the appearance of PIP structures (Fig. 3f, g), indicating that PSD-95 bridges the outer and inner phases. In order to explicate this phenomenon, we visualized the distribution of PSD-95 shared by GluN2B and STG that appeared at the interface between the inner and outer phases (shared PSD-95. Fig. 3h). In summary, PSD-95 played a role in the formation of the PIP structure by increasing the affinity between the two phases presumably by lowering their interfacial tension at the macroscopic level^6, 7, 8, 9, 11, 34^. The competitive binding-dependent phase transition between PIP and partial engulfment is also seen in the cases of RNA-binding proteins^9, 10^.

### PIP formation depends on linker length and multivalency of CaMKII

Next, we examined the role of the unique dodecameric structure of CaMKII in the PIP organization. Here, protein concentrations were fixed including the concentration of CaMKII-binding beads despite the change in the CaMKII valency. When the valency of CaMKII was decreased from 12 to 4, the PIP organization disappeared, and a homogenous condensate appeared instead (Fig. 4a, b). Decreasing the valency of CaMKII further to 2 caused the CaMKII to dissipate, and the remaining GluN2B, PSD-95, and STG formed a homogeneous condensate (Fig. 4c). Thus, high valency was found to be essential for the condensation, as demonstrated in a preceding computational study ^45^.

**Fig. 4.**
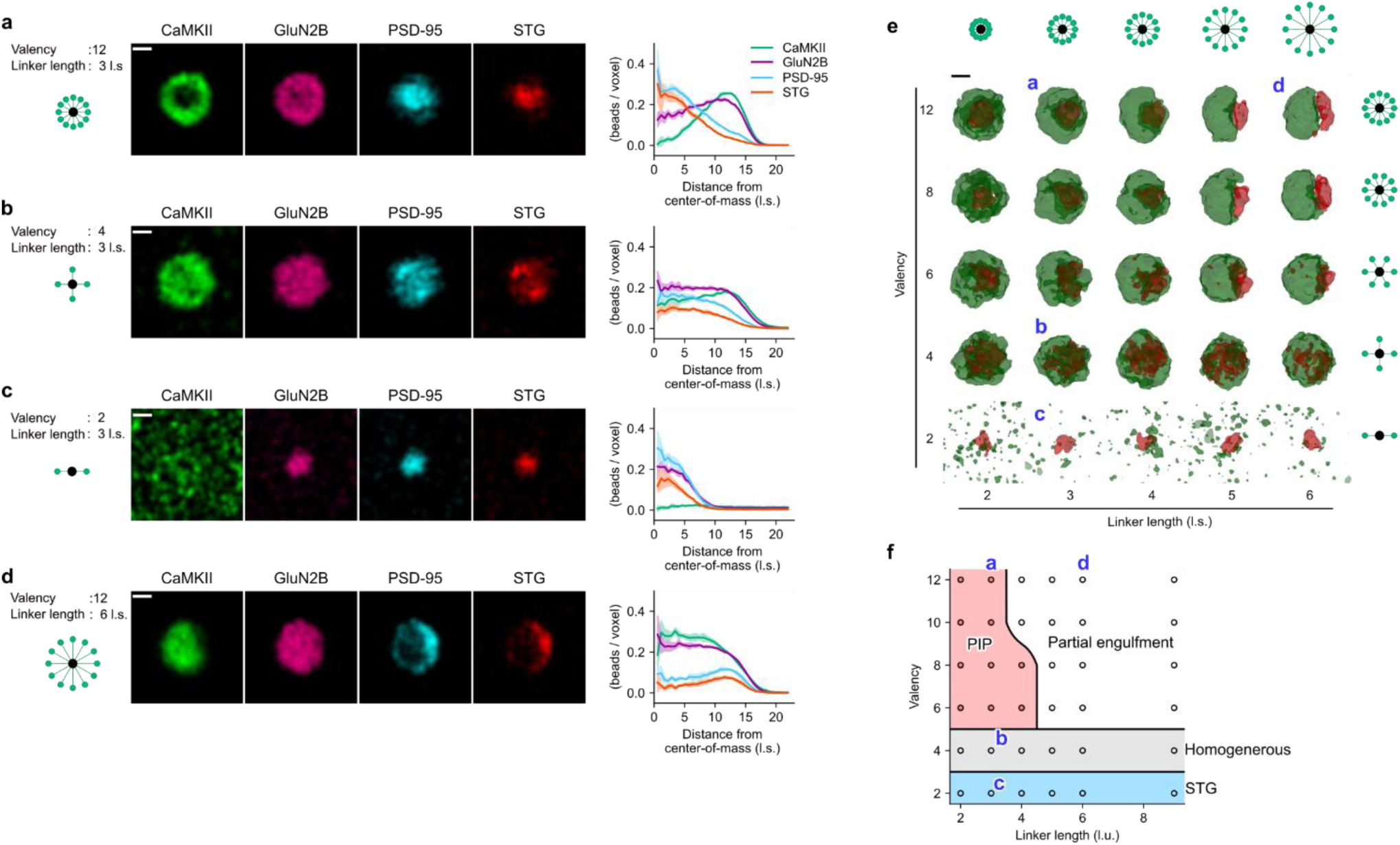
The PIP formation depends on both the valency and linker length of CaMKII. **a** The PIP formation under the standard CaMKII shape. CaMKII had 12 binding beads with a linker length of 3 l.s. Left: schematics; center: cross-sectional levels of constituent proteins; right: RDPs. **b** A homogeneous condensate formed if CaMKII had a valency of 4. **c** The homogeneous condensate of GluN2B–PSD-95–STG formed if CaMKII had a valency of 2. **d** A partially engulfed condensate formed if CaMKII had a linker length of 6 l.s. **e** The valency and linker-length dependence of two-phase condensates. Condensates in panels **a–d** are indicated. **f** Phase diagram of the LLPS. This phase diagram was determined based on Supplementary Fig. 4. Scale bar: 10 l.s.

We then tested the effect of the overall size of CaMKII protein by changing the linker length. When the CaMKII linker length was increased, two-phase condensates still appeared, but its shape was due to partial engulfment rather than PIP (Fig. 4d). This trend was consistently observed at a valency between 6 and 12, and the longer linker length favored the partial engulfment (Fig. 4e). The valency and linker-length dependence of CaMKII in LLPS is summarized in the phase diagram (Fig. 4f, Supplementary Fig. 3), which showed that the valency and linker length independently affected LLPS (Fig. 4f), although both had a significant effect on the LLPS without changing any other parameters, such as the concentration and binding energies.

### The shorter the CaMKII linker, the lower the surface tension of the condensate

Although the simultaneous binding of PSD-95 with GluN2B and STG was required for PIP in the case of concentration dependence (Figs. 2 and 3f–h), the ratio of PSD-95 bound to both GluN2B and STG did not change with the CaMKII linker length (Supplementary Fig. 5c), suggesting that another mechanism contributed to the phase transition between PIP and partial engulfment. The capillary theory demonstrates that both interfacial and surface tensions govern this type of phase transition^6, 7, 8, 9, 11, 34^. Here, through the cohesive force, we related the CaMKII linker length with the surface tension of the CaMKII–GluN2B phase.

Considering the difficulty of examining the surface tensions of the two-phase condensate, we simulated the binary mixtures of CaMKII and GluN2B and examined the spherical condensates (Fig. 5a, b; Supplementary Fig. 6a). We first found that the CaMKII hub with a longer linker length (6 l.s.) accumulated at the boundary region of the condensate than of the CaMKII hub with a short linker (3 l.s.) (Fig. 5a, b, top). The binding beads (kinase domain) of CaMKII at the border were not inwardly directed to the condensate center but radiated evenly from the center hub (Fig. 5a, b, bottom; Supplementary Fig. 6b). The radiation of binding beads may produce a surface tension under more realistic conditions.

**Fig. 5.**
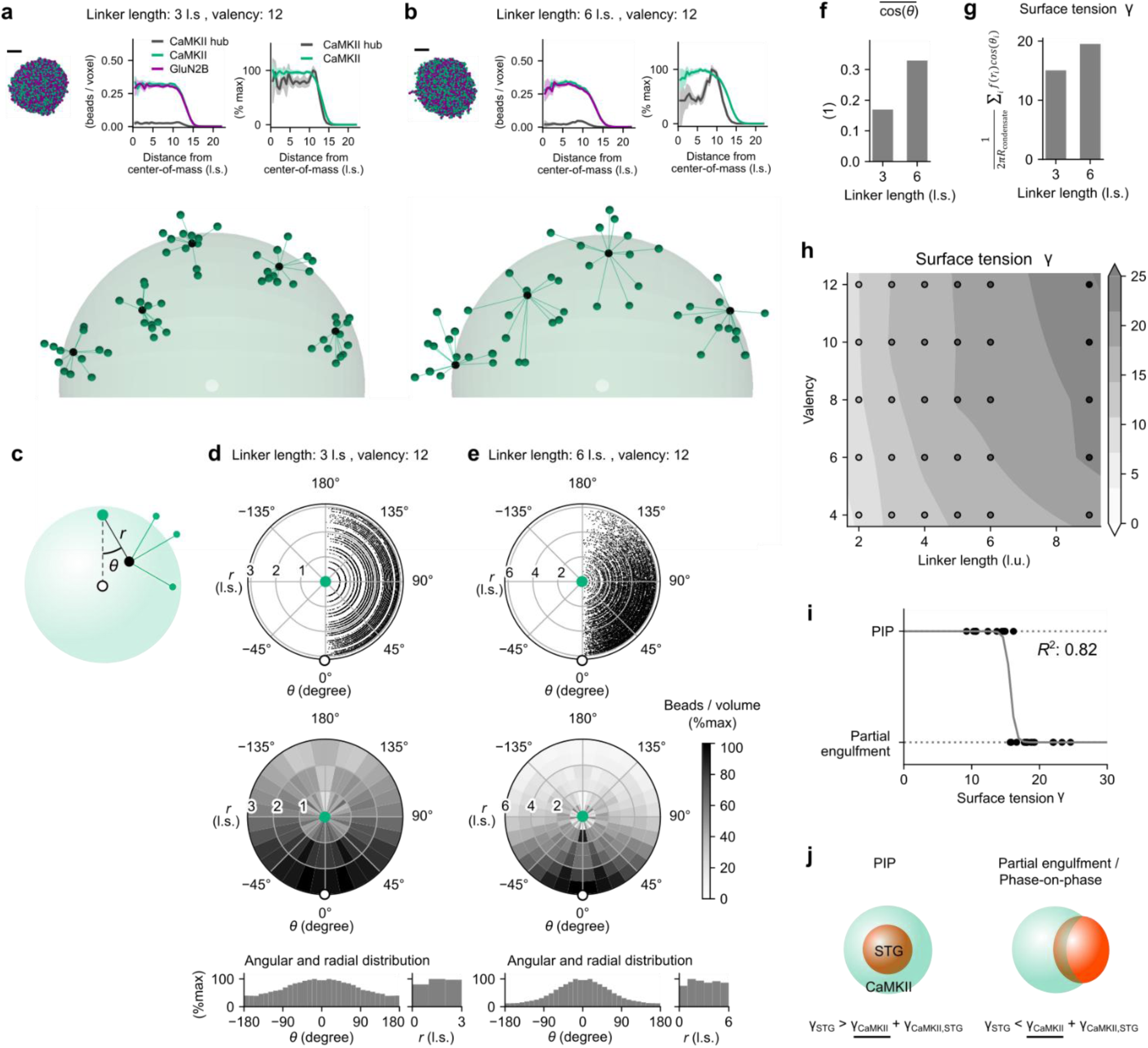
The analysis of CaMKII–GluN2B condensates explains the surface tension-dependent transition of a two-phase condensate. **a** The mixture of CaMKII and GluN2B forms a spherical-shaped condensate (left, top row), enabling numerical analyses of the putative surface tension. RDPs of CaMKII hub, CaMKII-binding beads (denoted as CaMKII), and GluN2B (center and right, top row) and the example sampling of CaMKII (bottom row) are also shown. Five CaMKII proteins were sampled to show spatial arrangements (condensate radius: 13.0 l.s.; Supplementary Fig. 6b, center). **b** Same as **a**, but the linker length of CaMKII was 6 l.s. Four CaMKII proteins were sampled in the bottom (condensate radius: 12.7 l.s.; Supplementary Fig. 6b, right). **c–e** Spatial distribution of CaMKII hub beads (black points in the left) when tethering binding beads were centered (green points); two standard cases are shown (**d, e**). Angles of the linker *i* versus the binding bead–condensate center axis (*θ_i_*) were obtained as well as distances between each CaMKII hub and binding bead (*r_i_*) (**c**); scatter plots showing the polar coordinates (top rows, in panels **d**, **e**). The polar (middle rows), angular (bottom rows, left), and radial distributions (bottom rows, right) are also shown. Levels in the polar distribution were normalized by the segmented volumes and those in the angular distribution were normalized by the segmented area. **f** Averaged cosine angles in the two cases. **g** Effective surface tension γ defined by ∑_*i*_ *f*(*r*_*i*_)cos (*θ*_*i*_)⁄2*πR*_condensate_ (Methods). **h** The surface tension γ in the plane of valency and linker length. **i** The surface tension γ explains the transition between PIP and partial engulfment. Efron’s *R*^2^ is displayed as a goodness-of-fit of the logistic regression. **j** Macroscopic determinant of the PIP and partial engulfment. The shape of the two-phase condensate is determined by the surface tension of the STG–PSD-95 phase, γ_STG_, and CaMKII–GluN2B phase, γ_CaMKII_, and their interfacial tension, γ_CaMKII,STG_. The calculated γ should be a part of γ_CaMKII_.

Therefore, we derived the putative inward force produced by the CaMKII-binding beads. In the current setup of MC simulation, the CaMKII linker had no tension associated with macroscopic properties. Nevertheless, the CaMKII binding beads could not move beyond a predetermined linker length from the center hub, and this spatial confinement can be considered as contraction force toward the center hub. This contraction force may partly be directed toward the center of the condensate and thus the inward cohesive force, whereas the tangential forces are canceled out under the spherical symmetry.

Based on this idea, we calculated the angle *θ_i_* between the linker *i* direction and the center-of-mass of a target condensate (Fig. 5c–e). *θ_i_* corresponds to the direction of linker-generated force with respect to the center of a condensate and showed a weak bias toward zero degrees if the CaMKII linker length was 3 l.s. (Fig. 5d, bottom). By contrast, if the CaMKII linker length was 6 l.s., the distribution of *θ_i_* was more concentrated at zero degrees (Fig. 5e). If the linker produces a contraction force, the component of this force directed to the center (cohesion force) can be calculated as cos(*θ*_*i*_). The averaged component, 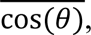 showed a clear difference between the linker lengths of 3 and 6 l.s. (Fig. 5f).

Effective contraction events of a linker occur when this is nearly fully stretched. Furthermore, if the total cohesion force toward the center is divided by the circumference length 2π*r*, this corresponds to the surface tension^46^. Therefore, we derived the putative surface tension γ and confirmed that the difference in 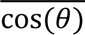 was preserved (Fig. 5f, g). In the plane of valency and linker length, the lower surface tension γ occupied the PIP region (Figs. 4f and 5h), the higher surface tension γ occupied the partial engulfment (Figs. 4f and 5h), and the surface tension γ explained this phase transition (*R*^2^ = 0.82, Fig. 5i). In the capillary theory, the difference between PIP and partial engulfment is determined by the surface tensions of two condensates γ_CaMKII_ and γ_STG_ as well as their interfacial tension γ_CaMKII,STG_^6, 7, 8, 9, 11, 34^. In the current case, the derived γ should constitute a part of γ_CaMKII_, although it did not correspond to the total γ_CaMKII_, including the surface tension due to GluN2B (Fig. 5j). The surface tension γ_CaMKII_ is expected to increase larger if the linker length was longer, which led to the phase transition from PIP (γ_STG_ > <u>γ_CaMKII_</u> + γ_CaMKII,STG_; Fig. 5j, left) to partial engulfment (γ_STG_ < <u>γ_CaMKII_</u> + γ_CaMKII,STG_; Fig. 5j, right).

### Modular networks in the CaMKII–GluN2B condensates

The simulation time required to form the stable CaMKII-GluN2B condensate was dependent on the shape of CaMKII. In order to quantify this time, we mimicked a fluorescence recovery after photobleaching (FRAP) assay on the CaMKII-GluN2B condensates (Methods, Fig. 6a–c). For this simulation, we used only local MC moves and considered the number of these as a proxy for time^37^. The recovery rate of CaMKII in the photobleached region was five-fold slower if the CaMKII had a short linker length (Fig. 6a, b). This slow rate was not derived from the longer lifetime of each CaMKII–GluN2B binding (Supplementary Fig. 7), the higher condensate concentration that may prevent the CaMKII movement (Supplementary Fig. 8), or the slow movement of independent CaMKII (Supplementary Fig. 9) but from their clustering connections (Fig. 6d–f). Herein, each GluN2B-mediated CaMKII–CaMKII connection (CaMKII–GluN2B–CaMKII connection) was considered as a unit CaMKII–CaMKII connection, and the connectivity of the simplified CaMKII network was quantified using the clustering coefficient (*CC*; Fig. 6d; Methods)^47^. The *CC* counts the number of connections between the neighboring CaMKII proteins to show the density of connections around the target CaMKII (Fig. 6d). The *CC* averaged over the target condensate explained the slowdown of CaMKII movement almost completely (*R*^2 =^ 0.80, Fig. 6c, e, f). The high *CC* presumably caused a higher rejection rate of local MC moves to produce the slow movement of CaMKII. Note that the MC moves differ from the real-time development as the MC simulation does not incorporate any diffusion rate constants or forward/backward-binding rate constants. Nevertheless, the cluster formation should also produce the slow diffusion of CaMKII in more realistic situations because the coupled CaMKII should have the larger Stokes radius. This has been demonstrated in MD simulation^48, 49^. The CaMKII with a short linker length and high valency is likely to show slow diffusion in the condensate due to its local clustering.

**Fig. 6.**
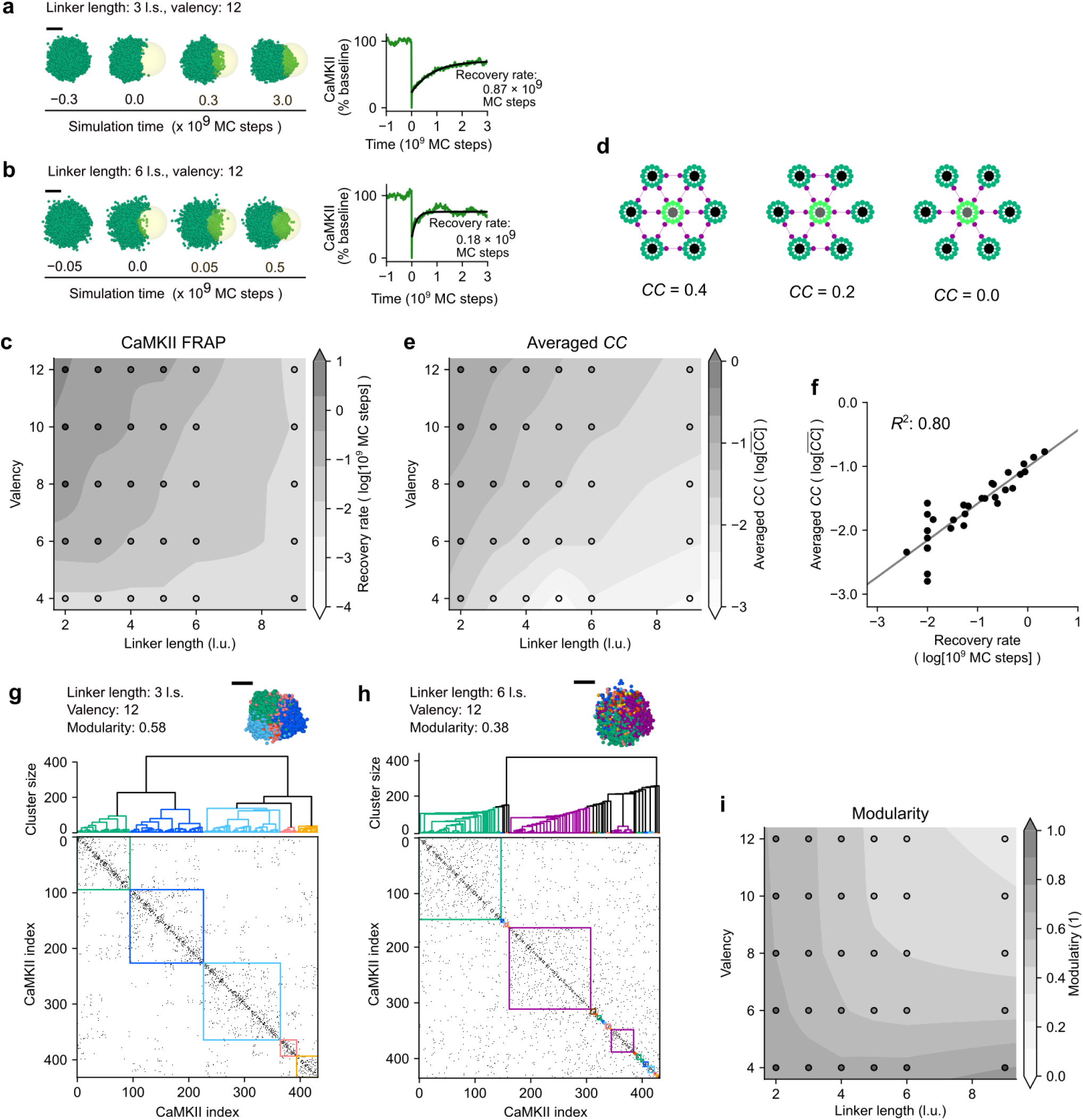
Recovery of CaMKII from the photobleaching correlates with the condensate network. **a** A FRAP experiment was mimicked on the CaMKII–GluN2B condensates (CaMKII linker length: 3 l.s., valency: 12; left). At the 0 MC step, a part of CaMKII condensate (yellow spherical region; radius: 8.7 l.s.) was made invisible, and then the recovery rate at this region was quantified (right). Scale bar, 10 l.s. **b** Same as **a**, but the linker length of CaMKII was 6 l.s. **c** Recovery rates in the plane of valency and linker length. **d** Clustering coefficient (*CC*) for the characterization of connectivity in CaMKII–GluN2B condensates (see Methods). In the condensate, CaMKII proteins (green) were bound to other CaMKII proteins via GluN2B (purple). If a target CaMKII protein (highlighted) binds to six neighboring CaMKII (*N* = 6) that have six surrounding connections (*n*_edge_ *=* 6, left case), this gives *CC =* 2*n*_edge_⁄*N*(*N* − 1) = 0.4. If *N* = 6 and *n_edge_ =* 3, this gives *CC =* 0.2 (center case), and if *N* = 6 and *n_edge_ =* 0, this gives *CC =* 0.0 (right case). In general, the larger *CC* denotes a denser local connectivity. **e** Averaged *CC* over all CaMKII proteins in the condensate (Methods). **f** The FRAP recovery rate is explained by the averaged *CC*. *R*^2^ is displayed as a goodness-of-fit of a linear function. **g** Adjacency matrix of CaMKII connections (CaMKII linker length: 3 l.s.; bottom). Each CaMKII connection via GluN2B is represented by a dot in the matrix. The order of CaMKII proteins was sorted using a hierarchical method for community detection (Girvan–Newman algorithm; Methods). Hierarchical communities are shown in the dendrogram (top), and clusters were colored below the threshold where 150 CaMKII proteins were assigned to the largest cluster. The shape of the condensate is shown (top, right). Scale bar, 10 l.s. **h** Same as **g**, but the linker length of CaMKII is 6 l.s. **i** The values of the measure “modularity” in the plane of valency and linker length (Methods).

The averaged *CC* only characterized the local connectivity. We further examined the more global characteristics of connectivity in the CaMKII–GluN2B condensates. In this analysis, we again targeted the GluN2B-mediated CaMKII connections and considered each CaMKII–GluN2B– CaMKII connection as a unit CaMKII–CaMKII connection (Fig. 6g, h). The simplified CaMKII network was analyzed using the Girvan–Newman algorithm, a hierarchical community detection method. The CaMKII connection exhibiting the highest edge-betweenness centrality was first selected and then removed. The repeats of this selection-removal cycle progressively decomposed the CaMKII network into multiple subnetworks whose CaMKII proteins were more densely connected (dendrograms in Fig. 6g, h). This decomposition revealed that CaMKII with the short linker had a more modular network, i.e., the CaMKII–GluN2B condensate comprised multiple subnetworks of several dozen CaMKII proteins (Fig. 6g). Conversely, CaMKII with the long linker had a smoother and more homogeneous network in the condensate (Fig. 6h). When the modularity was introduced as a measure of this global connectivity (Methods), the CaMKII–GluN2B condensates that had CaMKII with a short linker showed a large modularity if CaMKII had a valency >8 (Fig. 6i). If the CaMKII valency was <6 l.s., this trend disappeared presumably because the low valency CaMKII made many smaller subnetworks regardless of the linker length.

The modularity and averaged *CC* are different concepts (Fig. 6e, i): the modularity compares the relative number of intermodular connections against the background intramodular connections, whereas the *CC* quantifies the absolute degree of clustering in the network (Methods). The modular structure of CaMKII proteins always appeared when a short linker length was present, and the higher valency further slowed the movement of CaMKII. Such dynamic cluster formation and the associated slowdown of protein diffusion have been demonstrated in MD simulation^48, 49^.

## Discussion

Herein, we sought to elucidate the intrinsic factors that determine the topology of a two-phase condensate and developed a minimalistic computational model comprising four postsynaptic proteins. The quaternary mixture reproduced a form of two-phase condensates, termed PIP. The PIP did not always appear and depending on the concentrations of constituent proteins, the mixture showed homogeneous or partially engulfed condensation, the former of which was experimentally validated. Model analyses dissected the underlying physicochemical forces that drove these forms of condensates. The CaMKII PIP had a high valency and short linker length, whereas CaMKII exhibited a low surface tension, modular structure, and consequently slow diffusion.

Interestingly, those characteristics of CaMKII (i.e., low surface tension, modular structure, and slow diffusion) are aligned with the required functions of CaMKII as a synaptic memory unit. In general, LLPS shows poor stability^15, 16^. The continuing growth of condensates involves the absorption/sequestration of small condensates located in small domains, such as dendritic spines, which harbor the postsynaptic structures^50, 51^. Contrary to this process, synaptic condensates need to remain in place long enough to perform their functions. Theories for the coalescence process indicate that low surface tension and slow diffusion are both required for the long lifetime of small condensates^50, 52^. The shape characteristics of CaMKII lead to low surface tension and slow diffusion to ensure persistence for an extended period. The longer lifetime of the CaMKII condensates is important to activate downstream signaling for synaptic plasticity or synaptic tags^32, 53^. A recent study reported that a complex condensate actively regulates the condensate size^54^. In addition to PSD-95 anchoring the GluN2B-CaMKII condensate at PSD, the CaMKII shape might underlie the stability of small condensates in small postsynaptic domains.

For consistency with the preceding study^14^, we introduced GluN2B to form the CaMKII condensate. However, other client proteins, such as Shank3^55^, and Tiam1^56, 57^, are known to bind CaMKII together and may form CaMKII condensate. Also, CaMKII alone forms condensate at low pH^58, 59^. The characteristic shape of CaMKII is well conserved across metazoans^60^. The role of CaMKII condensate may not be restricted to the nanocolumn formation at PSD^14^ and the CaMKII condensate may play more universal roles in the formation of synaptic memory, thus synaptic plasticity.

The concentration dependence of LLPS revealed that the competitive binding of STG and GluN2B to PSD-95 was important for the PIP formation. Homogeneous condensation occurred at the high GluN2B concentrations (Fig. 2g, h) because PSD-95 was fully occupied with the excess amount of GluN2B, and the PSD-95–GluN2B complex further interacted with CaMKII to form the CaMKII– GluN2B–PSD-95 condensate (Supplementary Fig. 1d–f)^41^ while STG remained an association component. Similar competitive binding is also seen in the two-phasic separation in RNA and RNA-binding proteins^9, 10^. Thus, the competitive binding should be a motif of forming multiphase condensates.

To examine concentration dependence, the concentrations of STG and GluN2B were varied while those of CaMKII and PSD-95 were fixed. This was to demonstrate the quantification of the competitive binding of two client proteins toward PSD-95 and the binding of a client protein, GluN2B, to the scaffold protein, CaMKII. The concentration dependence of phase-separated structures on all four proteins may be a subject for a future study, although its explanation would be more complex.

The analyses of surface tension clarified that the directions of the CaMKII linkers were biased toward the center of condensate if the linker length was long (Fig. 5c–e). This was partly because the long linker length became closer in size to the condensate radius (Fig. 5a, b). The surface accumulation of CaMKII with a longer linker presumably minorly affected the surface tension. In a larger and open system, the surface accumulation of CaMKII would be a dominant factor with high surface tension because the summed cohesive force of CaMKII only at the surface region corresponds to the surface tension. Although the impact of the linker length on LLPS formation has been previously investigated^24, 26^, no study has yet provided systematic insights into how linker length controls the condensate shape. Herein, we successfully connected the microscopic linker length of CaMKII with the macroscopic forms of LLPS.

A recent study by Cai et al. showed that short linker CaMKIIα forms condensate together with GluN2B, but not long linker CaMKIIβ^44^. This clear trend was not observed in the current setup of simulation (e.g., Supplementary Fig. 6a). Presumably, this difference arises due to the different setups of the experiment. Unlike the experiment that we replicated^14^, Cai et al. used the full-length carboxyl tail of GluN2B, containing two CaMKII interaction sites^42^, and did not use the oligomeric form of GluN2B. Such difference in GluN2B might lead to the specific loss of CaMKIIβ condensate, although the mechanisms are yet unknown. The more detailed level of protein modeling might provide insight into the functional consequences of different CaMKII and GluN2B.

Our approach can easily be extended to incorporate many other postsynaptic proteins^5, 39, 61, 62,63^. Furthermore, our simulation can be extended to understand the other multiphasic LLPS that are based on divergent structures such as pentameric nucleophosmin (NPM1) in the nucleolus^6^, Rubbisco holo holoenzymes-EPYC1 condensate in pyrenoid^24^, and polycomb-repressive complex 1 (PRC1) family complexes in chromatin organization^28, 29^. Their unique forms would also be realized by the interfacial tensions of constituent molecules^6, 8, 9, 11^. The manipulation of the CaMKII linker length had drastic impacts on the multiphasic LLPS. Similar manipulations may be useful to control multiphasic LLPS in such systems.

## Methods

### *In vitro* phase separation experiment

Details of protein preparation are described elsewhere^14^. Briefly, PSD-95 with a thioredoxin tag and 3c protease site, CaMKIIα, DsRed2-spy catcher, spy-tag–carboxyl tail of STG (STGc), EqFP670-spy catcher, and spy-tag–carboxyl tail of GluN2B (GluN2Bc) were expressed in *Escherichia coli* strain BL21 DE3 RIL. These proteins were first purified via affinity chromatography using a Ni-NTA column (Nacalai Tesque). Tags were removed by cleavage with corresponding protease, and the purified DsRed-spy catcher or EqFP670-spy catcher were combined with spy-tag–STGc or spy-tag– GluN2Bc, respectively, to enable protein ligation. Combined proteins were further purified via size exclusion chromatography using a Hiload 16/60 Superdex 200 with ÄKTA GO FPLC system (Cytiva). CaMKII was further purified by the HiTrap Q column (Cytiva). CaMKII and PSD-95 were stained with iFluor488 and iFluor405, respectively, by using succinimidyl ester (AAT bioquest)^41^.

Purified proteins were combined in the observation buffer 50 mM Tris-HCl pH 8.0, 100 mM NaCl, and 1 mM TCEP in the presence of 20 μM calmodulin, 1 mM MgCl_2_, 1 mM ATP, and 1 mM CaCl_2_ to activate CaMKII. The final mixtures contained iFluor488-tagged CaMKII (CaMKII), eqFP670-tagged GluN2Bc (GluN2B), iFluor405-tagged PSD-95 (PSD-95), and DsRed2-tagged STGc (STG) at indicated concentrations. Protein concentration is denoted as monomer concentration throughout the study. For observation, 5 μL of the protein solution was injected into a sample chamber with a 15-mm diameter and 0.1-mm height. The sample was observed using a FV1500 confocal microscopy system (Olympus).

### LaSSI simulation

Simulations were performed on the LaSSI-simulation engine, which was developed for the MC simulation at coarse-graining levels^35^. All simulations were conducted in a 120 × 120 × 120 cubic lattice with periodic boundary conditions. The 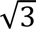 unit length of the cubic lattice was set as the unit length “lattice site” (l.s.) because this is used as the unit length of interactions and connections^35^. Various MC moves were implemented in LaSSI. The types and frequencies of MC moves used in the present study are described in Supplementary Table 1. Protein concentrations are described in Supplementary Table 2.

### Binding energy

Site-specific interactions were determined based on published binding affinities. We adopted −3.5, −5.25, and −3.0 *k_B_T* as binding energies between CaMKII and GluN2B^64, 65^, GluN2B and PSD-95^39, 41^, and PSD-95 and STG^39, 66^, respectively (Fig. 1b). Note, these shared similar energies because they had similar binding affinities.

IDR-mediated interactions were also determined based on the literature. In particular, the carboxyl tail of STG is rich in arginine residues, and a study reports that these produce a strong IDR-mediated affinity with PSD-95^41^. The carboxyl tail of GluN2B also contains arginine residues. We thus assigned values of −0.1, −0.11, −0.10, and −0.12 *k_B_T* as the affinities between STG and STG, STG and PSD-95, STG and CaMKII, and STG and GluN2B, respectively (Fig. 1c). Furthermore, all four proteins have electrically charged residues in their IDRs, which may cause IDR-mediated interactions. Thus, a small uniform affinity (−0.05 *k_B_T*) was assigned between all the beads as a baseline (Fig. 1c). The baseline affinity accounted for hydrophobicity and the other minor forces including the π–π, cation–anion, dipole–dipole, and π–cation interactions^7, 8^.

### Simulation time schedule

Each simulation began with random initial conditions, a system temperature of 1000 *T\**, and the following spherical constraining potential:

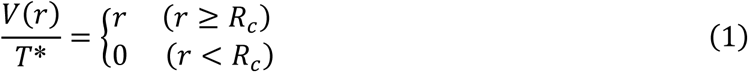

Where *r* denotes the distance from the center of the lattice space, 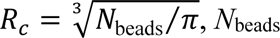 denotes the total number of beads, and *T\** denotes the unit of normalized temperature (*k_B_T*/*ε*)^35^. This constraining potential was introduced following preceding studies^7, 10^ to push all beads toward the center of the lattice space. The simulation ran over 5.0 × 10^7^ MC steps, then the system temperature was discontinuously decreased to 3.2 *T\** (Step 1, Supplementary Table 3). The simulation further ran over 1.0 × 10^8^ MC steps while decreasing the system temperature linearly from 3.2 to 1.0 *T\** or from 3.2 to 1.2 *T\** (Step 2, Supplementary Table 3). Then, the constraining potential was turned off, and the simulation ran over 1.0 × 10^11^ steps. The temperature was kept at 1.0 or 1.2 *T\** (Step 3, Supplementary Table 3). This time schedule enabled rapid convergence to the steady states.

Simulation results at the final time frame were collected for analysis. The last five time frames with an interval of 2 × 10^9^ MC steps were averaged to generate RDPs.

### Visualization

Steady-state profiles of mixtures were visualized using the following four methods. First, protein bead distributions in the 3D space were visualized using OVITO 3.10.2^67^. Front halves were removed, and only the rear halves are shown in Fig. 1 and Supplementary Figs. 1 and 9 for their visibility.

Second, protein beads were blurred by the Gaussian kernel with a standard deviation (SD) of 1.15 l.s., and their intensity levels are shown at the section that divided the center of the condensate (Figs. 1f, i, l, 2d–f, 3h, and 4a–d; Supplementary Fig. 1b, e, h). Intensity levels were normalized by the maximal level.

Third, RDPs from the condensate center-of-mass were generated according to previous literature^10, 35^. Briefly, the number histograms of each protein bead binning a unit sphere, *H*(*γ*_*n*_), were first calculated, then divided by the number histograms of lattice sites, *H*_*o*_(*γ*_*n*_) as follows:

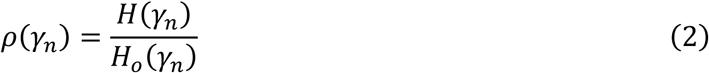

Where *γ_n_* denotes bins with a shell thickness of 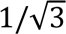 1.s, and *ρ*(*γ*_*n*_) gives the volume-occupation ratios of target proteins. The means and SDs were plotted based on data at five sampling time points with an interval of 2 × 10^9^ MC steps (Figs. 1g, j, m, 2d–f, 3h, and 4a–d; Supplementary Fig. 1c, f, i). This averaging was effective in compensating small sampling volumes in the small radius bins.

Fourth, the 3D shapes of the condensates were visualized using their rendered volumes (Figs. 2d–g and 4e; Supplementary Figs. 2, 4). Except for the volumes in Figs. 2d–f, all proteins were first rotated so that the barycenter of CaMKII beads was located on the left and that of PSD-95 beads was on the right. Then, the half maximal levels of blurred CaMKII and STG (Gaussian filter, *σ* = 1.15 l.s.) were used for the isosurface values for the marching cubes algorithm (Scikit-image 0.21.0)^68^, and obtained surface meshes were smoothed using the Humpy filter (*α* = 1.0 and *β* = 0.0, Trimesh 3.12.7)^69^. Smoothed meshes were visualized using Pyvista 0.43.3^70^ and used to determine the phase diagram of the condensates together with their sectioned intensity levels (Figs. 2 and 4; Supplementary Figs. 2 and 3).

### Surface tension

The MC sampling of LaSSI was intended to obtain the steady-state spatial distribution of beads, and linkers of LaSSI lack the tension associated with macroscopic characteristics of condensates. Nevertheless, a fully stretched linker prevents the tethered beads from being further apart, and in more realistic situations, this spatial constraint should produce tension. The summed tension directed to the center of condensate corresponds to the cohesion force, whereas the tangential forces are neutralized under the spherical symmetry. If the cohesion force is divided by 2*πr*, where *r* is the radius of the condensate, this forms a surface tension under the spherical symmetry condition^46^:

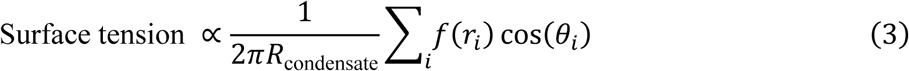

Where *r_i_* is the distance of *i*th CaMKII-binding bead from the hub bead, and *f*(*r*) is the relative strength of putative surface tension, which should have a positive value for almost fully stretched linkers. As an analogy to the local MC move of bead *i* being selected from the uniform distribution [−2, 2] and the MC move being rejected if *r_i_* exceeded the linker length^35^, *f*(*r*) was assumed as:

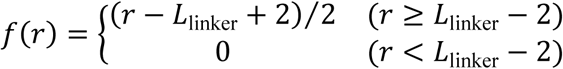

Where *L*_linker_ denotes the linker length.

### Connectivity analyses

The connections through site-specific interactions were examined using NetworkX 3.1. Each LAMMPS file from LaSSI was converted to an undirected multigraph, and the number of connected proteins were counted (Fig. 3a, c, and e).

### Graph theoretical analyses

We identified the CaMKII–GluN2B condensate as the largest connected network, (Figs. 5 and 6f–h)^37^. The connected network was further simplified into the graph that represented the connectivity of CaMKII proteins through GluN2B. In the CaMKII network, the *CC* for *i*th CaMKII protein, *CC_i_*, was calculated based on the following equation^36, 47^:

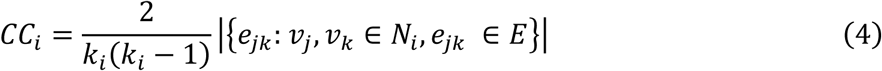

Where *N_i_* denotes the set of CaMKII bound to the *i*th CaMKII, *E* denotes the set of binding edges, *k_i_* denotes the number of the bound CaMKII, and *e_jk_* denotes the binding edge between *j*th CaMKII, *v_j_*, and *k*th CaMKII, *v_k_*. This corresponds to Fig. 6d, and intuitively, *CC_i_* denotes the number ratio of the formation of the triangular connection between CaMKII, one of which is the *i*th CaMKII. Then, the averaged 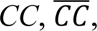 is defined as:

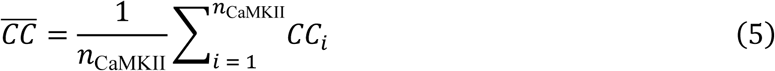

Where *n*_CaMKII_ denotes the number of CaMKII in the CaMKII network. The *CC* has a value in the range of [0, 1], and the higher value indicates a denser connectivity. We compared the averaged *CC* with the recovery rate *τ* from the photobleaching (Figs. 6c, e, f).

The overall characteristics of the CaMKII network were quantified using the Girvan– Newman algorithm for community detection (Figs. 6g–i) as described in the literature^71^. Here, the Girvan–Newman algorithm targets the CaMKII network and progressively removes edges (CaMKII– CaMKII connections) from this network. The removal was performed on the edge that showed the highest edge-betweenness centrality, i.e., the most important edge to form the connected condensate. The repeats of the removal split the CaMKII network into multiple connected networks. This process was depicted as a dendrogram (Fig. 6g, h).

To evaluate the global characteristics of the CaMKII network, we used the modularity, *Q*, as a measure of modular structure:

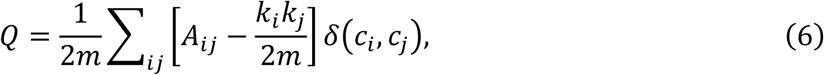

Where *m* is the total number of binding edges, *A* is the adjacency matrix (bottom in Fig. 6g, h), *k_i_* is the number of CaMKII bound to the *i*th CaMKII, *c_i_* denotes the community of the *i*th CaMKII, and *δ*(*i*, *j*) is the Kronecker delta. *Q* can be written as:

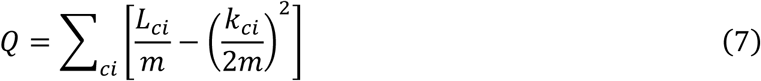

Where *ci* denotes the community index, and *L_ci_* is the number of intra-community links for community *ci*, *k_ci_* is the sum of the number of binding partners from all the CaMKII in community *ci*. In equation (7), the first term represents the number of intra-community connections, and the second term represents the total number of connections. Thus, *Q* represents the relative contribution of intra-community connections. *Q* has a value in the range of [−1/2, 1], and its higher value indicates a more modular network.

## Data availability

Raw data to reproduce figures, along with the Python files, have been deposited at https://github.com/urakubo/analyses_PIPS_camkii. Raw simulation data are available upon request.

## Code availability

All code used for modeling and simulation are available at https://github.com/pandeyv1990/modeling_and_simulation_PIPS_camkii. All code used to analyze data are available at https://github.com/urakubo/analyses_PIPS_camkii.

## Acknowledgments

We thank F. Dar and R. V. Pappu for providing us with the updated version of the LaSSI simulations engine. This project received funding from the Core Research for Evolutional Science and Technology (CREST), Japan Science and Technology Agency (JST) (Grant Number: JPMJCR20E4) to T.H., Y.H., and H.U. T.H. is supported by JSPS KAKENHI (JP19K06885), Kobayashi foundation and ISHIZUE2024 of Kyoto University. Y.H. is supported by Grant-in-Aid for Scientific Research JP18H05434, JP20K21462, and JP22K21353 from the MEXT, Japan, The Uehara Memorial Foundation, The Naito Foundation, Research Foundation for Opto-Science and Technology, Novartis Foundation, and The Takeda Science Foundation, HFSP Research Grant RGP0020/2019 and H.U. is supported by JSPS KAKENHI (JP24H02317, JP20K12062).

## Author contributions

V.P. and H.U. designed the research; V.P., H.U., and T.H. performed the research; V.P. and H.U. analyzed the data; H.U. and Y.H. supervised the research. All authors wrote the manuscript.

