## Supplementary Information for "The role of protein shape in multiphasic separation within condensates"

**Supplementary Table 1 | Types and frequencies of Monte Carlo (MC) moves**

| Move | Frequency | Frequency (FRAP simulation) |
|---|---|---|
| Local | 0.5445 | 1 |
| Multi-local | 0.001815 | 0 |
| Snake | 0.2722 | 0 |
| Translational | 0.0907 | 0 |
| Pivot | 0.0907 | 0 |

**Supplementary Table 2 | Protein concentrations**

| Composition (simulation) | CaMKII (beads/voxel) | GluN2B (beads/voxel) | PSD-95 (beads/voxel) | STG (beads/voxel) |
|---|---|---|---|---|
| CaMKII, GluN2B | 0.01 | 0.01 | | |
| PSD-95, STG | 0.01 | 0.01 | | |
| CaMKII, GluN2B, PSD-95 | 0.01 | 0.01 | 0.01 | |
| GluN2B, PSD-95, STG | 0.01 | 0.01 | 0.01 | |
| CaMKII, GluN2B, PSD-95, STG | 0.01 | 0.01 | 0.005 | 0.005 |
| CaMKII, GluN2B (Surface tension simulation) | 0.01 | 0.01 | | |
| CaMKII, GluN2B (FRAP simulation) | 0.003 | 0.01 | | |

**Supplementary Table 3 | Simulation time schedule**

| Mixtures | Simulation steps | | | | |
|---|---|---|---|---|---|
| | Step 1 | Step 2 | Step 3 | Step 4 | Step5 |
| Binary | $5 \times 10^7$ MC steps<br>1000 to 3.2 $T$* | $1 \times 10^8$ MC steps<br>3.2 to 1.0 $T$* | $1 \times 10^{11}$ MC steps<br>1.0 $T$* | | |
| Ternary | $5 \times 10^7$ MC steps<br>1000 to 3.2 $T$* | $1 \times 10^8$ MC steps<br>3.2 to 1.0 $T$* | $1 \times 10^{11}$ MC steps<br>1.0 $T$* | | |
| Quaternary | $5 \times 10^7$ MC steps<br>1000 to 3.2 $T$* | $1 \times 10^8$ MC steps<br>3.2 to 1.2 $T$* | $1 \times 10^{11}$ MC steps<br>1.0 $T$* | | |
| Binary for the simulation of surface tension and FRAP | $5 \times 10^7$ MC steps<br>1000 to 3.2 $T$* | $1 \times 10^8$ MC steps<br>3.2 to 1.2 $T$* | $1 \times 10^{11}$ MC steps<br>1.2 $T$* | $1 \times 10^8$ MC steps<br>1.2 $T$*<br>only for observation | |
| Supplementary Video 1 | $5 \times 10^7$ MC steps<br>1000 to 3.2 $T$* | $1 \times 10^8$ MC steps<br>3.2 to 1.0 $T$* | $1 \times 10^{11}$ MC steps<br>1.0 $T$* | $1 \times 10^{11}$ MC steps<br>1.0 to 1.2 $T$*,<br>CaMKII activated | $1 \times 10^{12}$ MC steps<br>1.2 $T$* |

**Supplementary Video 1 | Activation of CaMKII results in the PIP formation.**

**Simulation time schedule:** First, inactive CaMKII, GluN2B, STG, and PSD-95 were mixed where the inactive CaMKII was designed to not bind any other proteins. The simulation ran standardized steps 1, 2, and 3 to reach an equilibrium state (Supplementary Table 3). After these steps, we observed a STG–PSD-95–GluN2B condensate and CaMKII in the diluted phase (Supplementary Fig. 1g–i). CaMKII was then activated and the simulation continued until the next equilibrium state (steps 4 and 5 in Supplementary Table 3). During step 4, the system temperature was linearly increased from 1.0 $T^*$ to 1.2 $T^*$ for rapid convergence. The video shows the transition of the condensate during steps 4 and 5.

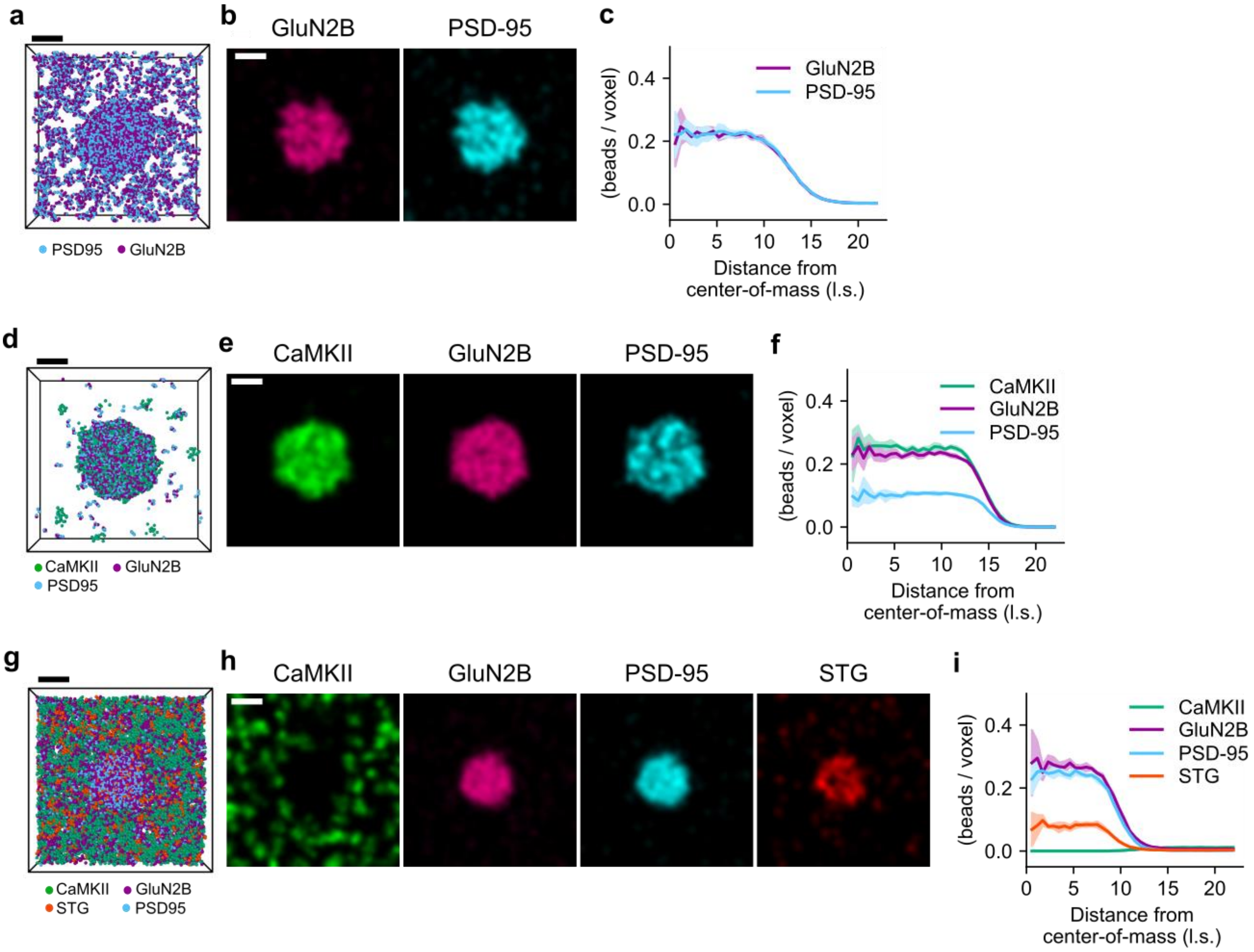


**Supplementary Fig. 1 | Simulated condensates are consistent with experimentally observed condensates.**

**a–c** The binary mixture of CaMKII and GluN2B formed a homogeneous condensate. Panels show snapshots of their spatial distribution (**a**), cross-sectional distribution (3D Gaussian-filtered, $\sigma = 2$ l.s.). (**b**), and averages and standard deviations of radial distribution profiles (RDP; $n = 5$) (**c**), which were consistent with results in previous studies[1, 2]. **d–f** The ternary mixture of CaMKII, GluN2B, and PSD-95 formed a homogeneous condensate, which is consistent with results in a prior study[1]. **g–i** The mixture of inactivated CaMKII, GluN2B, STG, and PSD-95 formed a homogeneous condensate, consistent with previous results[3]. Snapshots in panels **a**, **d**, and **g** show only the back half of the protein distribution. Scale bar: 10 l.s.

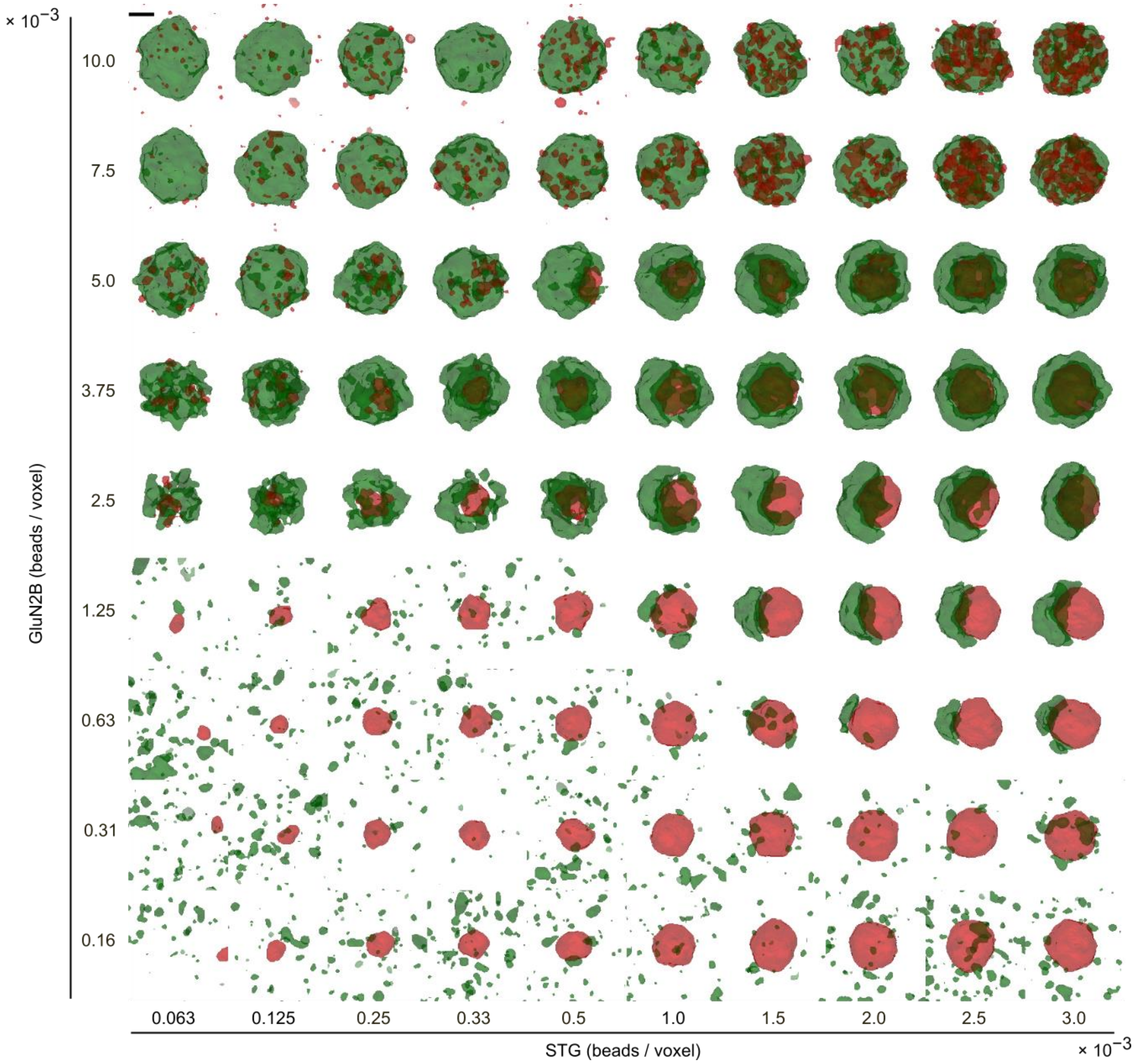


**Supplementary Fig. 2 | Rendered shapes of two-phase condensates of CaMKII–GluN2B and STG–PSD-95 under the concentration plane of STG and GluN2B.**

CaMKII–GluN2B condensates are colored green, and STG–PSD-95 condensates are colored red. The phase diagram in Figure 2h was created based on the visual inspection of this Figure as well as sectional density images. Scale bar: 10 l.s.

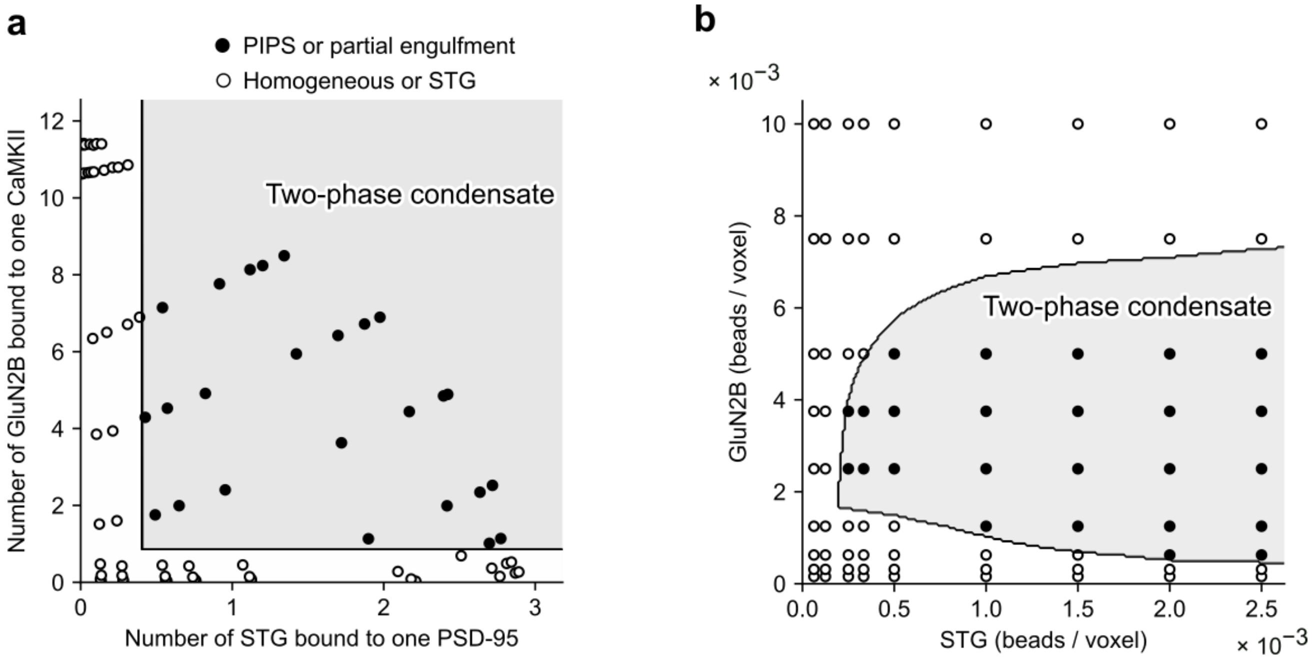


**Supplementary Fig. 3 | The two-phase condensate is characterized by two types of protein binding.**

**a** Appearance of one- or two-phase condensates (open and closed circles, respectively) in the plane of the average number of GluN2B proteins bound to one CaMKII and the average number of STG proteins bound to one PSD-95. A simple decision tree (depth: 3, solid lines) gave the region of two-phase condensates with 100% accuracy (gray-shaded area); thresholds were 0.85 for the GluN2B–CaMKII binding and 0.41 for the STG–PSD-95 binding. **b** The one- or two-phase condensates in the concentration plane of STG and GluN2B. This panel is identical to Figure 3e.

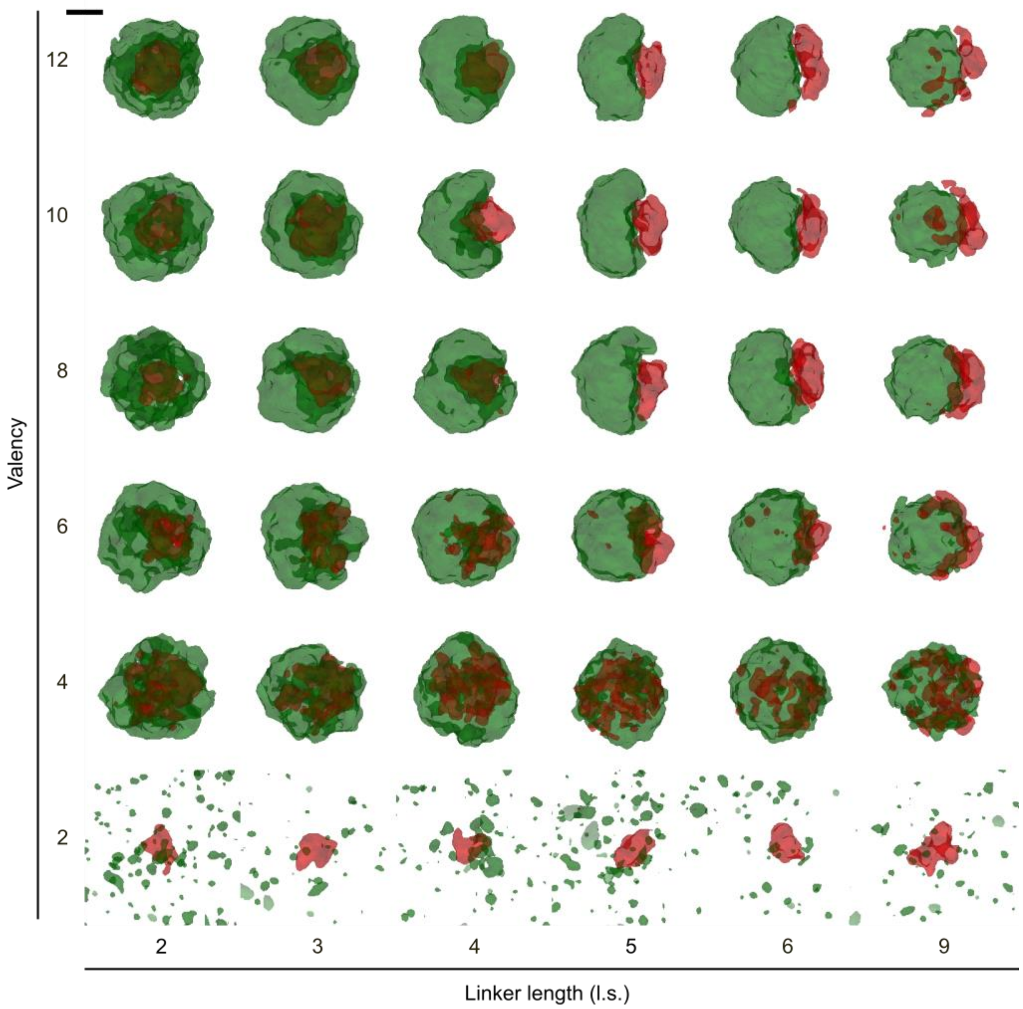


**Supplementary Fig. 4 | Rendered shapes of condensates under various valencies and linker lengths of CaMKII.**

The CaMKII phase is green, and the STG phase is red. The phase diagram in Figure 4f was created based on the visual inspection of this figure as well as sectional density images. Scale bar: 10 l.s.

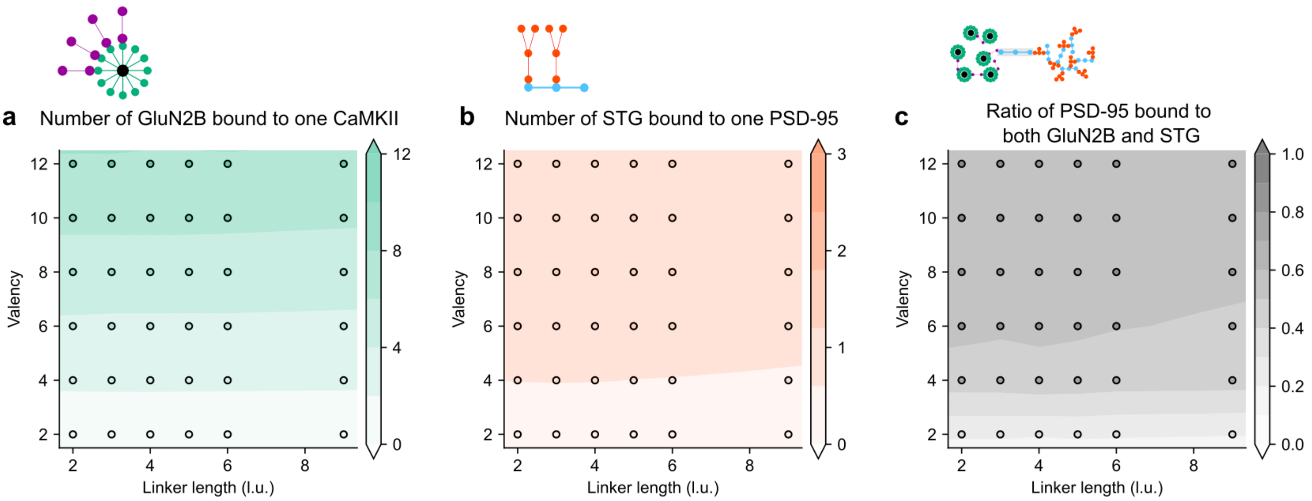


**Supplementary Fig. 5 | The number ratio of competitive binding of GluN2B and STG to PSD-95 does not differ depending on the linker length of CaMKII.**

**a** Average number of GluN2B bound to one CaMKII in the plane of CaMKII linker length and valency. **b** Average number of STG bound to one PSD-95. **c** Number ratios of PSD-95 bound to both GluN2B and STG.

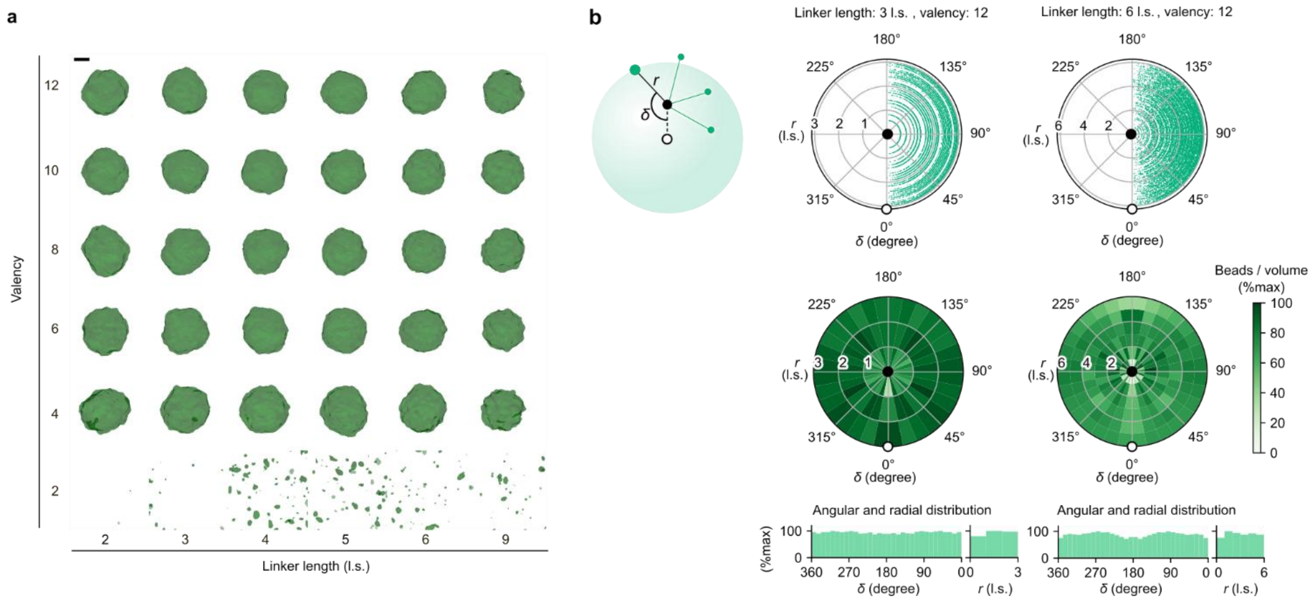


**Supplementary Fig. 6 | The CaMKII–GluN2B mixture forms spherical-shaped homogeneous condensate regardless of the shape of CaMKII, and the condensate CaMKII shows the spherically uniform distribution of CaMKII-binding beads.**

**a** Rendered shapes of the CaMKII–GluN2B condensates in the plane of valency and linker length. The mixtures of GluN2B and CaMKII underwent homogeneous condensation if CaMKII had a valency >4. Scale bar: 10 l.s. **b** The spatial distribution of CaMKII-binding beads (green points in the left panel). Two standard cases are shown. Angles of each binding bead from the axis of the CaMKII hub-condensate center ($\delta$) and distances from the CaMKII hub ($r$, left) were obtained; scatter plots show the polar coordinates (top row). The polar distribution (middle row) and angular and radial distribution (bottom row) are shown. Each segmented level in the polar distribution was normalized by the segmented volume and each segmented level in the angular distribution was normalized by the segmented area.

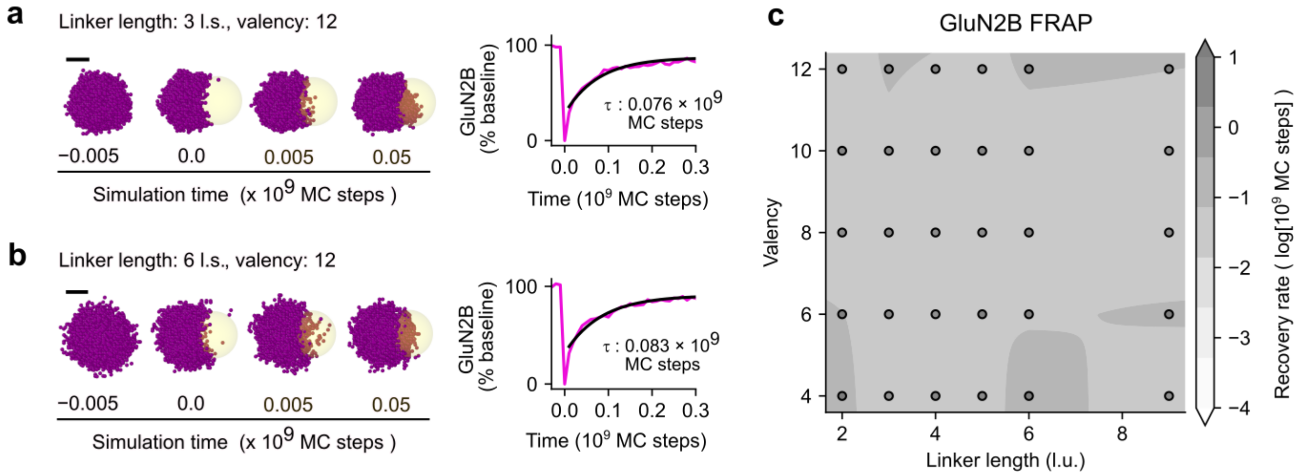


**Supplementary Fig. 7 | GluN2B consistently showed a similar recovery rate from photobleaching regardless of the shape of CaMKII.**

**a** Photobleaching was applied to a spherical area (radius: 8.7 l.s.; yellow area at 0 MC step; left), and the recovery rate of GluN2B at the bleached region (yellow areas) was quantified (τ, right). Scale bar, 10 l.s. **b** Same as **a**, but the linker length of CaMKII was 6 l.s. **c** Recovery rates of GluN2B in the plane of CaMKII linker length and valency.

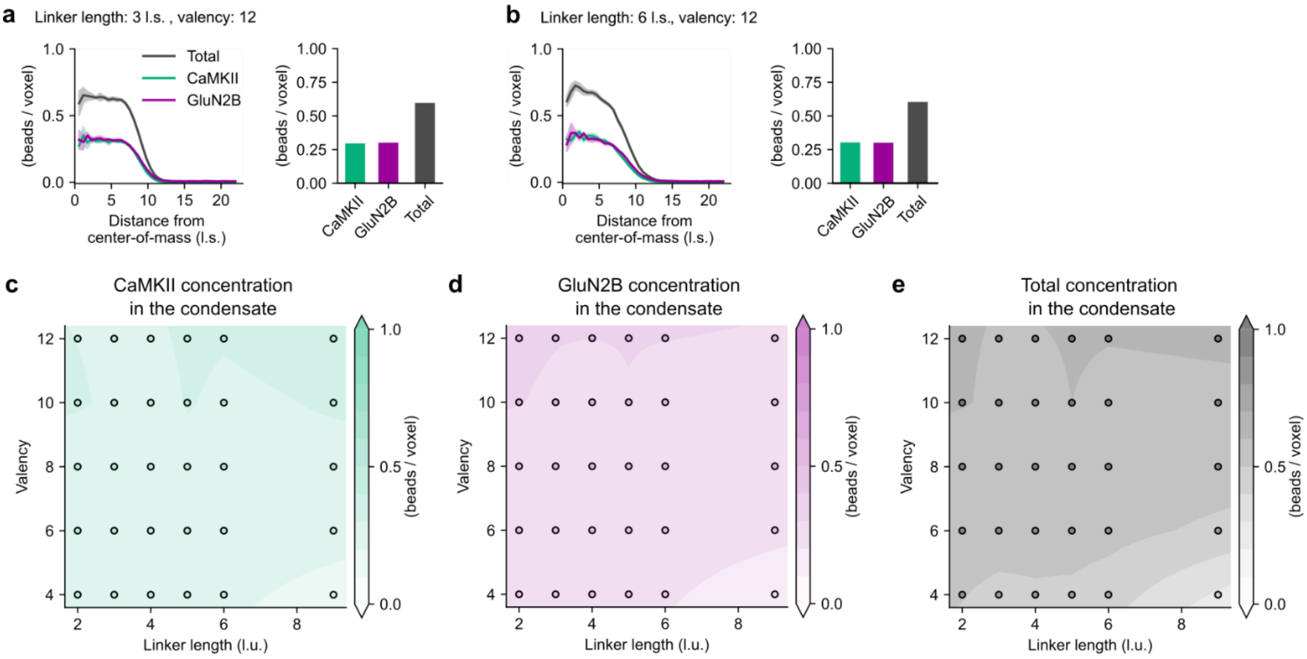


**Supplementary Fig. 8 | The shape of CaMKII does not affect the condensate concentrations of CaMKII and GluN2B.**

**a** RDPs for the CaMKII–GluN2B condensate (linker length: 3 l.s., valency: 12; left panel), and condensate concentrations of CaMKII and GluN2B (right). Each condensate region for the concentrations was defined as the region over the half-maximal levels of blurred beads of CaMKII (Gaussian filter, $\sigma$ = 1.15 l.s.). **b** Same as **a**, but the linker length of CaMKII was 6 l.s. **c–e** Concentrations of CaMKII (**c**), GluN2B (**d**), and the total bead concentration (**e**) in the plane of linker length and valency.

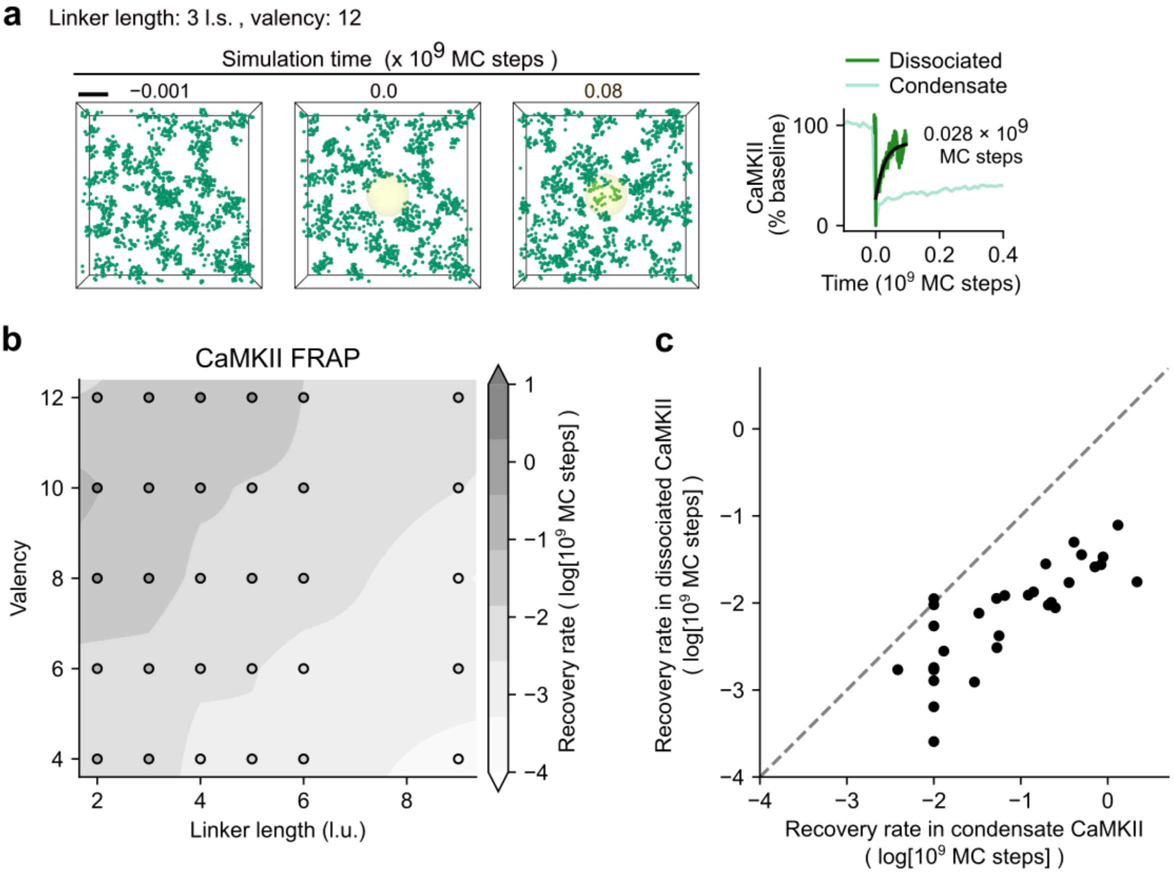


**Supplementary Fig. 9 | Slow recovery rates from photobleaching in condensates were not due to the slow movement of dissociated CaMKII.**

**a** Photobleaching was applied to dissociated CaMKII (linker length: 3 l.s., valency: 12; left panel), and the recovery rate was quantified (black line in the right panel; 0.028 × $10^9$ MC steps). The recovery profile was the average of eight simulations and was 30-fold faster than that in the CaMKII–GluN2B condensate (light green line; 0.87 × $10^9$ MC steps in Fig. 6a). Scale bar: 10 l.s. **b**, Recovery rates in dissociated CaMKII in the plane of linker length and valency. **c**, The dissociated CaMKII had ~10-fold faster recovery rates than the condensate CaMKII; therefore, the diffusion of dissociated CaMKII was not the time-limiting process. Points were mapped from data points in Figure 6h and panel **b**.
